# Kidney failure alters parathyroid Pin1 phosphorylation and parathyroid hormone mRNA binding proteins leading to secondary hyperparathyroidism

**DOI:** 10.1101/2021.12.06.470227

**Authors:** Alia Hasan, Yael E. Pollak, Rachel Kilav-Levin, Justin Silver, Nir London, Morris Nechama, Iddo Z. Ben-Dov, Tally Naveh-Many

## Abstract

Parathyroid hormone (PTH) regulates calcium metabolism and bone strength. Chronic kidney disease leads to secondary hyperparathyroidism (SHP) which increases morbidity and mortality. In experimental SHP, the increased *PTH* gene expression is due to enhanced *PTH* mRNA stability, mediated by changes in its interaction with stabilizing AUF1 and destabilizing KSRP. Pin1 isomerizes target proteins and leads to KSRP dephosphorylation. In SHP, Pin1 isomerase activity is decreased and phosphorylated KSRP fails to bind *PTH* mRNA, resulting in high *PTH* mRNA stability and levels. However, the up- and down-stream mechanisms by which kidney failure stimulates the parathyroid remain elusive. We now reveal a pathway where kidney failure induces parathyroid Pin1 phosphorylation, linking Pin1, KSRP and PTH mRNA stability as key players for the high PTH expression in SHP. We identified by mass-spectrometry, changes in rat parathyroid proteome and phosphoproteome profiles induced by impaired renal function, including KSRP phosphorylation at Pin1 target sites. Furthermore, both acute and chronic kidney failure led to parathyroid-specific Pin1 Ser16 and Ser71 phosphorylation, which disrupts Pin1 activity. Accordingly, pharmacologic Pin1 inhibition, that mimics the decreased Pin1 activity in SHP, increased PTH expression ex-vivo in parathyroid organ cultures and in transfected cells, through the *PTH* mRNA protein-interacting element and KSRP phosphorylation at potential Pin1-binding motifs. Therefore, kidney failure leads to loss of parathyroid Pin1 activity by inducing Pin1 phosphorylation. This predisposes parathyroids to increase PTH production through impaired *PTH* mRNA decay that is dependent on KSRP phosphorylation. Pin1 and KSRP phosphorylation and the Pin1-KSRP-*PTH* mRNA axis thus drive SHP.

## Introduction

Parathyroid hormone (PTH) is central to calcium homeostasis and bone strength. Changes in extracellular calcium are sensed by the parathyroid calcium sensing receptor (Casr), which in turn regulates PTH secretion and synthesis (1-4). Secondary hyperparathyroidism (SHP) is common in chronic kidney disease (CKD) and when poorly controlled, results in increased incidence of bone fractures, vascular calcification and mortality (5-7). SHP also occurs in acute kidney injury (AKI), because of abrupt deterioration in kidney function, and is frequently complicated by other mineral metabolism disorders (8-10). We have previously shown in experimental models of SHP due to dietary induced hypocalcemia or uremia, high PTH serum and mRNA levels that are mainly due to post-transcriptional upregulation. The increase in *PTH* mRNA stability and steady-state levels correlates with changes in the binding of *trans* acting proteins, AUF1 (AU rich binding factor 1) and KSRP (K-homology splicing regulatory protein) to a defined evolutionary conserved adenine/uridine (AU) rich element (ARE) in the *PTH* mRNA 3’-untranslated region (3’-UTR) (11). The peptidyl prolyl *cis/trans* isomerase (PPIase) Pin1 coordinates these mRNA-protein interactions by regulating phosphorylation induced protein conformation (12).

AUF1 (encoded by *HNRNPD*) recognizes ARE sequences in mRNA 3’-UTRs inducing either target mRNA stability or decay (13-15). KSRP promotes rapid decay of ARE-containing RNAs. KSRP contains four contiguous K homology (KH) domains that recognize AREs and mediate RNA binding, mRNA decay, and interactions with the exosome and poly(A)-specific ribonuclease (PARN) (16). KSRP phosphorylation links KSRP-mediated mRNA degradation to extracellular signals (17). p38 MAPK-mediated KSRP Thr692 phosphorylation impairs KSRP-RNA interactions and increases abundance of a variety of target mRNAs (17-19). We have previously identified an additional KSRP phosphorylation site and Pin1 target at Ser181, which prevents KSRP mediated *PTH* mRNA degradation (12). Ser181 KSRP phosphorylation and Pin1 activity were also reported in the PTH-induced rapid decay of the type IIa sodium-phosphate cotransporter (*Npt2*) mRNA in renal opossum kidney cells (20).

Pin1 binds phosphorylated Ser/Thr residues followed by proline (pSer/Thr-Pro) in target proteins (21). This interaction catalyzes the *cis-trans* isomerization of the peptide bond, thereby changing the activity, stability and localization of target proteins participating in many cellular functions, including transcription and post-transcriptional regulation of gene expression (21-24). Pin1 interacts with several RNA binding proteins, among them KSRP, AUF1, HuR, Histone stem-loop-binding protein (SLBP) and cytoplasmic polyadenylation element binding protein (CPEB), that regulate mRNA decay, as well as other aspects of mRNA lifecycle (12, 25-28). These interactions depend on cell type and environmental stimuli. We have previously shown that Pin1 interacts with KSRP in parathyroid glands and in human embryonic kidney (HEK) 293 cells (12). Pin1-mediated KSRP Ser181 dephosphorylation favors KSRP-*PTH* mRNA 3’-UTR binding and thus *PTH* mRNA decay. In SHP, parathyroid Pin1 isomerase activity is decreased and as a consequence phosphorylated Ser181 KSRP fails to induce *PTH* mRNA decay, resulting in increased PTH mRNA and serum levels. Mice with genetic deletion of Pin1 (*Pin*^*-/-*^ mice) had a 3-fold increase in serum PTH levels compared to wild type (wt) littermates, with no change in serum calcium and phosphorus levels. The *Pin*^*-/-*^ mice also had a 2-fold increase in PTH mRNA and protein content in the parathyroid glands, in agreement with their high serum PTH levels. These results support the central role for Pin1 in determining basal PTH levels in vivo (12, 29).

Pin1 is composed of an N-terminal WW protein interaction domain and a C-terminal catalytic PPIase domain (21, 30-32). Pin1 binding and catalytic activity is tightly controlled by several post-translational modifications. Protein kinase A (PKA)-mediated Pin1 serine (Ser)16 phosphorylation abolishes the interaction of Pin1 with its substrates (33). Death-associated protein kinase 1 (DAPK1) phosphorylates Pin1 at Ser71 in the catalytic active site and inhibits its isomerase activity (34, 35). Therefore, Pin1 Ser16 or Ser71 phosphorylation is central to Pin1 inactivation. In addition, Pin1 cysteine (Cys)113 residue is essential for Pin1 catalytic activity, and its oxidation inhibits Pin1 cellular function to promote tau and amyloid precursor protein turnover in neurons (36). Mutagenesis of Cys113 to alanine (Ala) results in loss of Pin1 protein activity (32). Screening an electrophilic fragment library, targeting Cys113, identified Sulfopin as a highly selective Pin1 inhibitor directed towards Pin1’s active site. Sulfopin exhibits potent target engagement and phenocopies *Pin*^*-/-*^ mice (37).

The molecular mechanisms that regulate Pin1 activity towards its target proteins are not fully understood. In particular, it is not clear what steers the decrease in Pin1 activity leading to high PTH levels in SHP (12, 38). We now show that experimental CKD alters overall parathyroid protein expression and phosphorylation. Kidney failure leads to parathyroid specific Pin1 phosphorylation that is consistent with the decreased Pin1 isomerase activity and the resulting parathyroid KSRP phosphorylation at Pin1 target motifs. Pharmacologic Pin1 inhibition, that mimics the decreased Pin1 activity in kidney failure, increases *PTH* expression in parathyroid glands in culture and in transfected cells in a manner that is dependent on KSRP phosphorylation and the PTH mRNA *cis* acting element. We suggest that CKD-induced Pin1 and hence KSRP phosphorylation and the Pin1-KSRP-PTH mRNA axis are central to the increased *PTH* expression of SHP.

## Results

### Chronic kidney disease induces changes in parathyroid proteome and phosphoproteome composition

To characterize molecular changes in chronic kidney disease (CKD) induced secondary hyperparathyroidism (SHP), we performed the first proteome and phosphoproteome analysis of the minute micro-dissected parathyroid tissue from kidney failure and control rats. To induce kidney failure and SHP, male rats were fed either a control or an adenine-rich high phosphorus diet for 2 weeks that led to the expected increase in serum creatinine, phosphate and PTH levels (not shown) (12). At the end of the diet period, the two parathyroid glands of each rat were micro-dissected from surrounding thyroid tissue and pools of 2 rats were analyzed by mass spectrometry (MS) in triplicates. MS proteome analysis identified 4950 proteins groups, encoded by 3790 genes (several identifiers per gene often arising from splice isoforms). Parathyroid tissues from kidney failure rats could be distinguished from controls based on the overall proteome profiles (**Fig 1A**). Specifically, 173 and 110 proteins were up- and down-regulated, respectively, in extracts obtained from kidney failure induced SHP glands compared to control glands (**Fig 1B** and **Supplemental Table 1**). These changes in parathyroid gland protein expression highlight potentially modified pathways in CKD (**Supplemental Fig 1**), leading to SHP. As can be expected, hyperparathyroidism (hyperplastic state) was accompanied by changes in ribosome protein levels, protein translation, mRNA stability and secretion pathways (**Fig 1C**). Proteome analysis thus revealed moderate yet significant profile alteration in SHP and put forward pathways for future studies.

**Figure 1.**
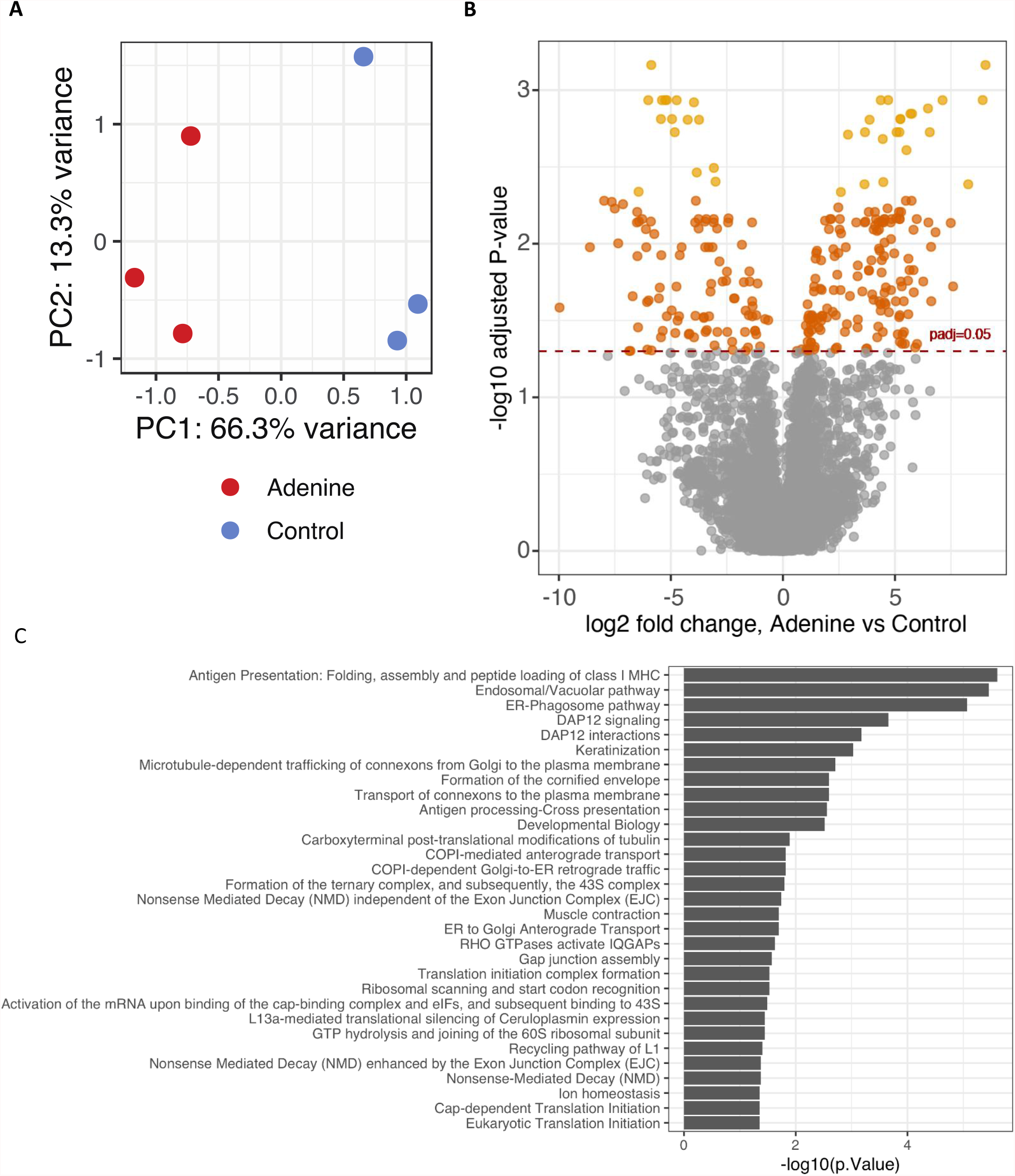
Experimental chronic kidney disease leads to proteomic changes in parathyroid tissue. Rats were fed either a control or an adenine-rich high phosphorus diet for 2 wk to induce kidney failure and extracts were prepared from micro-dissected parathyroid glands for MS proteome analysis. **(A)** Principal component analysis (PCA) plot of mass spectrometry intensity data (n=4950 protein groups) showing proteomics-based distinction of adenine (red) from control (blue) glands. **(B)** Volcano plot showing the landscape of protein dysregulation in adenine glands. Brown light brown dots represent proteins at adjusted p-value cutoffs <0.05 and 0.005 respectively. Positive Log2 fold change represents proteins that are increased in kidney failure parathyroids. **(C)** Enrichment of Reactome pathway-associated proteins among the differentially expressed parathyroid proteins (see also Figure S1).

Phosphoproteome analysis of pools of parathyroid extracts from 6 control and 6 kidney failure rats recovered 5370 parathyroid phosphopeptides [87.3% pSerine (Ser), 12.0% pThreonine (Thr) and 0.7% pTyrosine (Tyr)], encompassing 3884 phosphosites in 2222 phosphoproteins. Results of biological theme or function enrichment analysis of the identified phosphoproteins, with the detectable proteome as background using DAVID, are shown in Supplemental Fig 2A. Protein phosphorylation and kinase activity and structure terms dominate the enrichment lists, and transcription regulation is also prominent. This result is consistent with previous studies (39) demonstrating that regulatory proteins, such as transcription factors and kinases, are more often subjected to post-translational regulation via phosphorylation than are metabolic enzymes. Enrichment of ErbB and mTOR signaling-related proteins is also apparent and warrants further studies in this model. The enrichment for mTOR signaling is consistent with our previous findings showing that mTOR activation is central to the development of SHP and parathyroid cell proliferation induced by either uremia or hypocalcemia in rats (40). There was also a significant increase in CLOCK protein phosphorylation in kidney failure rat parathyroids (P < 0.0005) at a conserved site that alters CLOCK nuclear localization and hence function (Supplemental Table 2). This is supported by the recent findings showing that a circadian clock operates in parathyroid glands and that CLOCK and downstream cell cycle regulators are disturbed in uremia and may contribute to parathyroid proliferation in SHP (41).

We then examined up- and downregulation of phosphosites and phosphoproteins. The detected phosphoproteome was subjected to phosphosite- and gene-centered enrichment analyses by means of a contemporary approach, ‘PhosR’ (42). To this end, we processed an additional pool of parathyroid glands from adenine-rich high phosphorus diet fed kidney failure rats and from control rats, and only phosphopeptides identified in both runs were included in the subsequent analysis (n=4171, Supplemental Table 2). Phosphosite dysregulation in adenine compared to control rats is shown in **Fig 2A**. The median log_2_ fold change of phosphosite abundance was 0.14 (∼10.5% increase), significantly higher than 0 (no change) according to a one-sample Wilcoxon test (p<0.00001). Consistent with this shift, there were 460 significantly upregulated phosphosites (11.0%) and only 348 significantly downregulated sites (7.4%), indicating a possible global increase in protein phosphorylation. This may be expected in light of increased intracellular phosphate concentration in uremic SHP (43). However, we observed downregulation of the sodium-dependent phosphate transporter, Pit1, encoded by *Slc20a1* (log2 fold change -1.17, p<0.01), and respectively reduced Pit1;Ser269 and Pit1;Ser271 phosphorylation. In this regard, possible increases in Mapk1/3 (Erk1/1) activity, proposed to transduce the downstream signaling of high phosphate levels sensed by unknown mechanisms (that may include Pit1 and the calcium sensing receptor, Casr) (44) is suggested by enrichment of a potential MAPK phosphorylation motif (45, 46) <PLpSP> among the recovered phosphosites in the adenine (kidney failure) pools (median log_2_ fold change 0.22, ∼16.3% increase) (Supplemental Fig 2B). Thus, overall the phosphoproteome analysis suggests a globally increased phosphorylation state.

**Figure 2.**
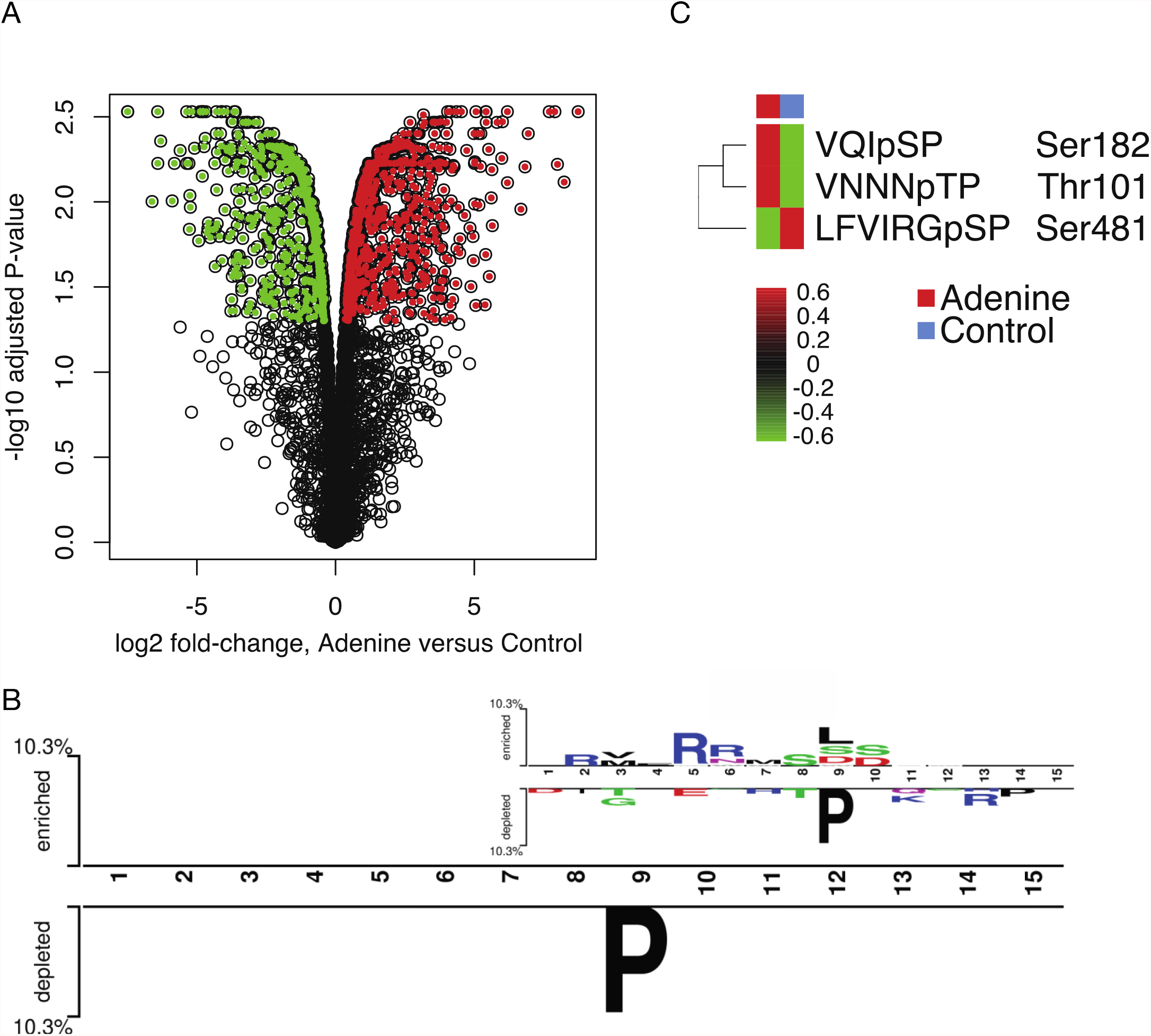
Experimental chronic kidney disease leads to phosphoproteomic changes in parathyroid tissue. (**A**) Volcano plot showing phosphosite dysregulation in parathyroid extracts from rats fed adenine-rich (2 pools of 6 rats) versus control diet (2 pools). Colored dots represent phosphorylation sites (red, increased; green, decreased) at adjusted p-value <0.05. Positive Log2 fold change represents phosphoproteins that are increased in kidney failure parathyroids. The top upregulated phosphosites (log2 fold change ≥ 5) were Sptbn1:S2099, Bckdk:T32, Hist3h2ba:S5, Tmem69:S13, Hp1bp3:T51 and Virma:S1578, while Casr:S1066, Evl:S333, Lmo7:S619, Lpp:T630, Krt86:S47 and Ptbp1:S140 were the most downregulated sites. (**B**) Two sample (48) presenting the only statistically robust sequence difference between phosphosites increased in adenine rats and phosphosites decreased in adenine rats; the odds ratio for P in the 9th position in adenine vs. control was 0.79, Bonferroni-corrected p<0.0001. The inset shows additional differences for which the Bonferroni-corrected p-values were insignificant. (**C**) KSRP phosphorylation sites detected by mass spectrometry analysis as above. Peptides that are potential target sites for Pin1, Ser182 (VQIS, valine-glutamine-isoleucine-serine-proline), Thr101 (VNNNT, valine-asparagine-asparagine-asparagine-threonine-proline) were increased and Ser481 (LFVIRGSP, leucine-phenylalanine-valine-isoleucine-arginine-glycine-serine-proline) was decreased in parathyroids obtained from adenine (kidney failure) rats.

PhosR prioritizes potential kinases that could be responsible for the phosphorylation change of phosphosites based on kinase recognition motif and phosphoproteome dynamics (42). Kinase activity scores that sum changes in phosphorylation of targets, for which the targeting kinase is prioritized with a probability score, pointed to significantly increased activity of specific kinases in adenine (kidney failure) parathyroids (**Supplemental Fig 2B**). Among others, there was increased activity of calcium/calmodulin-dependent protein kinase type II (Camk2a), Prkg1 (cGMP-dependent protein kinase 1) and PKA (Prkaca). Phosphorylated Camk2a has been shown to block PTH secretion ex-vivo and negatively correlate with serum calcium levels in patients with primary hyperparathyroidism, and thus may act to oppose hypersecretion (47), but studies in SHP are lacking. Numerous protein targets for Prkg1 phosphorylation are implicated in modulating cellular calcium levels. cAMP/PKA (Prkaca)-dependent mechanism inactivate the isomerase Pin1 by phosphorylating Pin1 Ser16 in the WW domain, leading to dissociation of Pin1 from its substrates (33). PKA activated Pin1 Ser16 phosphorylation could lead to the parathyroid Pin1 decreased activity in SHP induced by either experimental CKD or prolonged hypocalcemia (12).

Next, applying the “Two Sample Logo” approach (48) we found global differences between adenine fed kidney failure rat parathyroid proteins with enriched and depleted phosphosites (**Fig 2B**). Proline (Pro) in the position after phosphorylated Ser, Thr or Tyr was markedly depleted in glands from adenine fed kidney failure rats (odds ratio 0.79, Bonferroni-corrected p-value <0.0001). Phosphorylated Ser/Thr-Pro motifs are potential targets for Pin1 that often leads to conformational changes that induce decreased phosphorylation of these sites in target proteins. Our current findings of reduced pSer/pThr followed by Pro imply fewer potential Pin1 targets in kidney failure rat parathyroid extracts that could be due to changes in kinase activity, leaving Pin1 with less potential targets. The inset shows additional, less robust findings from this analysis (Fig 2B). In light of the latter result and our prior findings of decreased Pin1 activity in kidney failure parathyroids (12), we narrowed our search for proteins that were phosphorylated on Ser/Thr-Pro motifs as potential targets for Pin1 isomerase activity. We have previously shown that the *PTH* mRNA decay promoting protein KSRP is a target protein for Pin1, that when over-expressed induces KSRP dephosphorylation at Ser181 and increased *PTH* mRNA stability and levels. Pin1 knock-down or inhibition had the opposite effect, to increase KSRP phosphorylation and *PTH* mRNA levels in transfected cells (12). Indeed, experimental CKD that decreased Pin1 activity (12), led to parathyroid KSRP hyperphosphorylation at three Pin1 potential target sites. KSRP phosphorylation at Ser182 (Ser181 in human) and Thr101 were both increased (p=0.016, Fig 2C). KSRP pSer481, another yet uncharacterized potential Pin1 target, was decreased in parathyroids obtained from kidney failure rats (p=0.041, Fig 2C and Supplemental Table 2). These KSRP post-translational modifications may be due to the decreased Pin1 activity in SHP parathyroids (12) and would alter KSRP-*PTH* mRNA interaction and contribute to the increased *PTH* mRNA levels in SHP. We therefore studied the changes that affect Pin1 isomerase activity in kidney failure induced SHP and their effect on KSRP mediated *PTH* expression.

### Chronic and acute kidney failure induce parathyroid Pin1 specific phosphorylation that parallels decreased Pin1 activity in secondary hyperparathyroidism

Pin1 isomerase activity is regulated by phosphorylation at Ser16 and Ser71 that disrupts its interaction with target proteins and catalytic isomerase activity respectively (35). Because parathyroid Pin1 isomerase activity is decreased in CKD (12), we set out to determine if CKD affects parathyroid Pin1 phosphorylation. For that, we performed an in vitro kinase assay using parathyroid extracts from control and kidney failure induced SHP rats and recombinant GST-Pin1. Rats were fed a control or an adenine-rich high phosphorus diet as above (12). Extracts from pools of micro-dissected parathyroid glands from control and CKD rats were incubated with recombinant GST-Pin1 or GST as control and [γ-^32^P]ATP, and analyzed after GST-pulldown by SDS-PAGE and autoradiography. SHP rat parathyroid extracts were more potent in inducing GST-Pin1 phosphorylation, compared to control extracts (Fig 3A&B). Therefore, experimental CKD leads to increased parathyroid phosphorylation activity towards Pin1.

**Figure 3.**
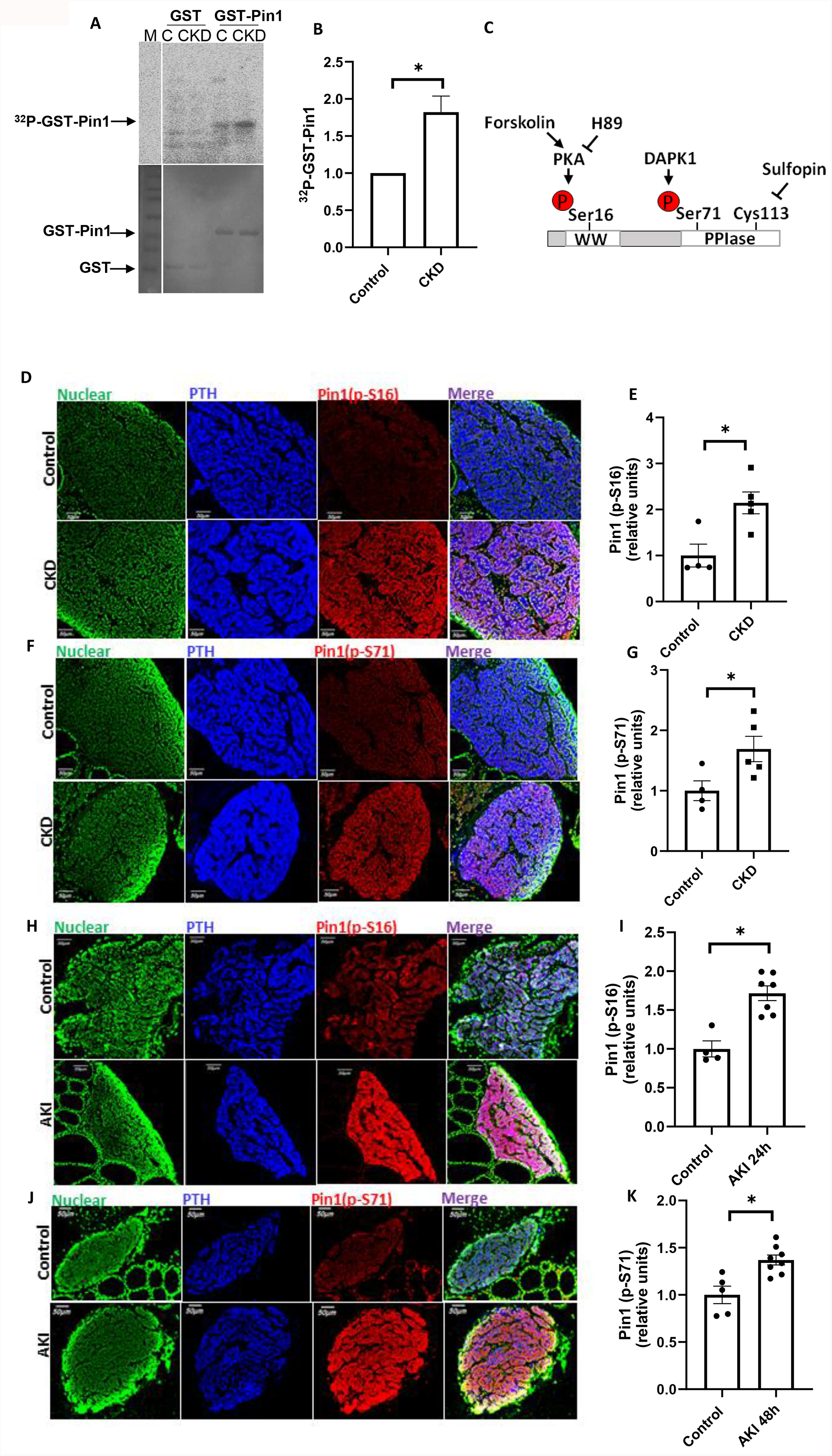
Experimental chronic kidney disease leads to increased parathyroid Pin1 Ser16 and Ser 71 phosphorylation. **(A**) In vitro phosphorylation assay of recombinant GST-Pin1 by parathyroid extracts from control (**C**) and experimental chronic kidney failure (CKD) rats. Male rats were fed either an adenine-rich high phosphorus or a control diet for 2 wk to induce kidney failure and SHP. Extracts from pools of 11-13 rats in each group were incubated with GST or GST-Pin1 and [γ^32^P] ATP. ^32^P-labeled GST-Pin1 was detected by SDS PAGE autoradiography (top) and Coomassie blue-staining for input GST and GST-Pin1 recombinant proteins (bottom). (**B**) Quantification of the intensity in autographs as in **A** and data from additional 3 repeat experiments. (**C)** Schematic representation of Pin1 protein showing the WW protein-binding and the PPIase catalytic (PPIase) domains. The regulatory Ser16 and Ser71 phosphorylation sites and their respective kinases as well as the Cys113 site and pharmacological modulators are shown. (**D-G)** Pin1 immunofluorescent staining of thyroparathyroid sections from control and kidney failure (CKD) rats as in A. Sections were stained for SYTOX nuclear staining (green) and antibodies for PTH (blue), Pin1 p-Ser16 (**D**) and p-Ser71 (**B**) (red) and merge (purple). (**E&G)** Quantification of relative fluorescence intensity for Pin1 in slides as in D and F and additional rats, measured by Image J software. (**H-K**) Pin1 immunofluorescent staining of thyroparathyroid sections from control and folic acid induced acute kidney injury (AKI) mice at 24 h (**H**) or 48 h (**J**) post folic acid or vehicle (control) injection. Sections were stained as above with Pin1 p-Ser16 (**H**) or p-Ser71 (**J**). (**I&K**) Quantification of Pin1 staining in H and J and additional mouse samples. Results in B, E,G,I,K are presented as mean±SE of fold change, compared to controls. *, p<0.05.

We then examined parathyroid specific Pin1 Ser16 or 71 phosphorylation (Fig 3C), that disrupts Pin1 isomerase activity (35). We could not detect these Pin1 pSer16 and pSer71 containing peptides in our MS analysis, possibly due to limitations of the trypsin protein digestion method (49). Immunofluorescence (IF) analysis of thyro-parathyroid sections from kidney failure and control rats using specific targeted pSer16 and pSer71 antibodies (34) showed increased phosphorylation of both Pin1 Ser16 and 71 in kidney failure SHP parathyroid glands, compared to glands from control rats (Fig 3D-G). This increase in Ser16 and Ser71 phosphorylation was not associated with detectable changes in total Pin1 protein levels, as expected (Supplemental Fig 3) (12).

To study Pin1 phosphorylation in an additional model of SHP, we induced acute kidney injury (AKI) by a single injection of high dose folic acid in mice (9, 10). Folic acid led to the expected increase in serum urea and PTH levels at 24 and 48 h (Supplemental Fig 4) (9, 50). IF analysis showed that AKI mice had increased parathyroid Pin1 pSer16 and pSer71 phosphorylation compared to vehicle-injected mice (Fig 3H-K). As in kidney failure rat parathyroids, there was no change in total Pin1 levels (not shown). Therefore, SHP is characterized by increased parathyroid phosphorylation activity towards Pin1. Specifically, Pin1 Ser16 and 71 phosphorylation is increased in two experimental models of chronic (CKD) and acute (AKI) SHP. The increased Ser16 and 71 phosphorylation correlates with the previously described decreased Pin1 activity and high *PTH* expression in SHP (12).

### PKA activation induces Pin1 Ser16 phosphorylation and mediates the increase in *PTH* expression in parathyroid organ cultures and in transfected cells

PKA, which is more active in uremic parathyroid glands (Supplemental Fig 2B), phosphorylates Pin1 Ser16 in the WW domain, leading to Pin1 dissociation from its substrate (33) (Fig 3C). To determine if alterations in Pin1 phosphorylation affect PTH expression, we pharmacologically manipulated Pin1 in parathyroid glands ex-vivo and in transfected cells. We first studies the direct effect of PKA activation on PTH secretion ex-vivo in mouse thyro-parathyroid glands in organ culture, as there is no functional parathyroid cell line (51, 52). The specific PKA activator forskolin increased PTH accumulated in the culture media at 1 and 3 hours post incubation, compared to vehicle treated cells (Fig 4A). Therefore, PKA activation increases PTH secretion from parathyroid glands in culture.

**Figure 4.**
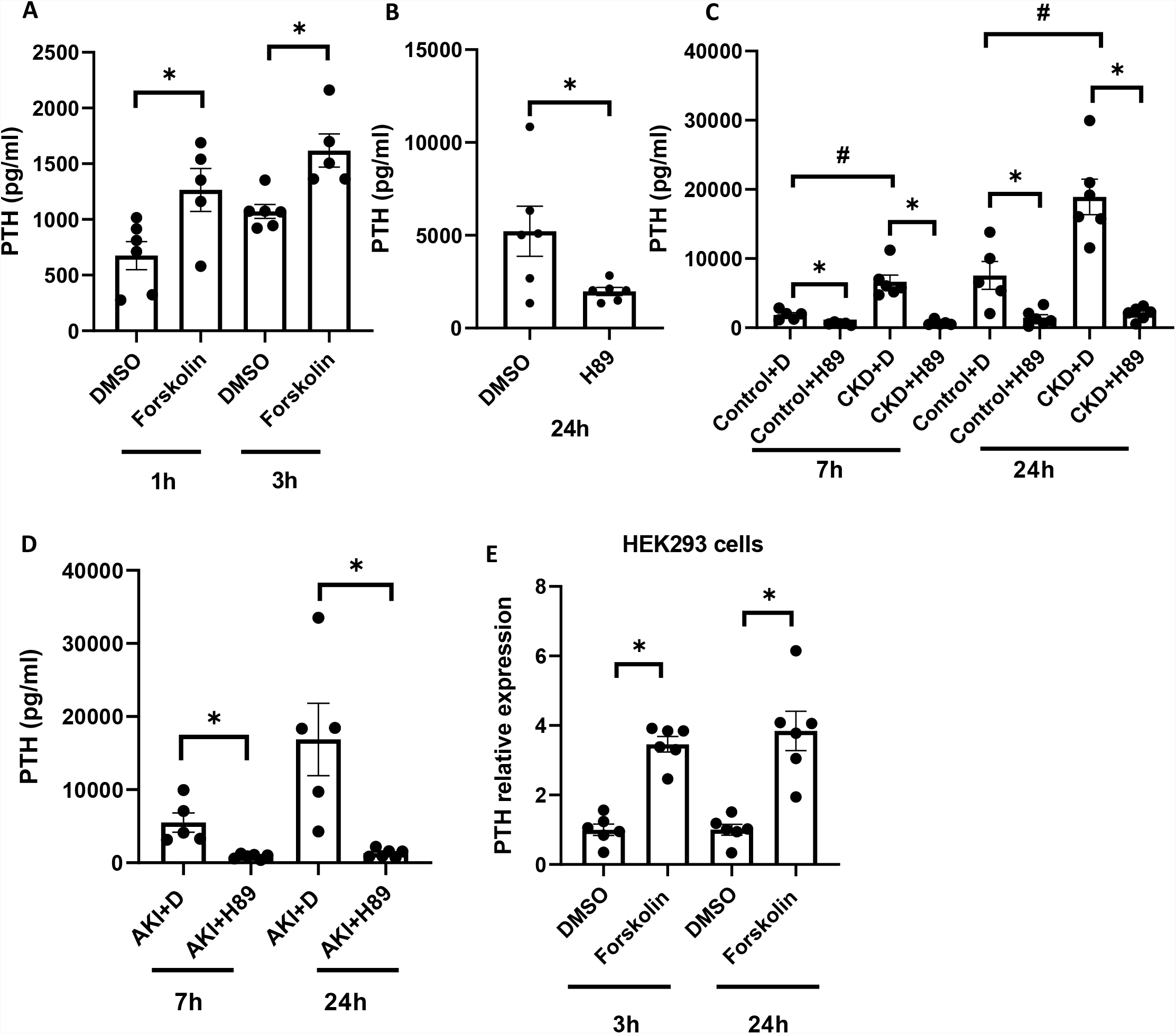
PKA induced Ser16 phosphorylation at the Pin1 WW domain increases PTH expression in mouse parathyroid glands in culture and in transfected HEK293 cells. (**A**) Mouse thyro-parathyroid glands were microdissected and the two glands form each mouse maintained in culture with the PKA activator forskolin (100 µM) or vehicle (DMSO). PTH accumulated in the culture medium was measured at 1 and 3 h. (**B**) Rat micro-dissected parathyroid glands (two glands form each rat) were maintained in culture with the PKA inhibitor H89 (150 µM) or vehicle (DMSO) and secreted rat PTH measure at 24 h. (**C**) Mice were fed a normal (Control) or an adenine-rich high phosphorus diet to induce renal failure (CKD). At 2 wk, thyro-parathyroid glands were cultured with H89 or DMSO (+D) as in B. PTH accumulated in the culture medium was measured at 7 and 24 h. (**D**) Mice received a single injection of folic acid (240 mg/kg) to induce acute kidney injury (AKI). At 24 h thyro-parathyroid glands were removed and cultured with H89 or DMSO (+D) and PTH accumulated in the culture medium measured. (**E**) HEK293 cells were transiently transfected with an expression plasmid for the human PTH gene. Transfected cells were incubated with forskolin (10 µM) or DMSO. PTH mRNA levels at 3 and 24 h, normalized to GAPDH, were determined by qRT-PCR. Results are presented as mean±SE of accumulated PTH (**A-D**) or fold change of PTH mRNA (E), compared to vehicle treated glands or cells. *, p<0.05 compared to control animals or cells incubated with DMSO; #, p<0.05 compared to control+D.

We then studied the effect of PKA inhibition on PTH secretion. The specific PKA inhibitor, H89 (N-[2-p-bromocinnamylamino-ethyl]-5-isoquinolinesulphonamide) led to a decrease in secreted PTH compared to vehicle (DMSO) in rat micro-dissected parathyroid glands in culture (Fig 4B). We also added H89 to mouse thyro-parathyroids from control and adenine-rich high phosphorus induced kidney failure mice (Fig 4C). Of interest, thyro-parathyroids from CKD mice cultured in control medium for 7 and 24 h, secreted more PTH compared to glands from mice with normal renal function, indicating that stimulation of the parathyroid by kidney failure in vivo is preserved in vitro (Fig 4C)(51). Importantly, H89 decreased PTH secretion from parathyroids in culture of both control and kidney failure mice (Fig 4C). The effect of PKA inhibition to decrease PTH secretion was also evident in folic acid-induced AKI mouse thyro-parathyroid glands in culture (Fig 4D), similar to its effect on parathyroid glands form normal renal function and kidney failure mice and normal rats (Fig 4B-C). Therefore, pharmacological PKA activation and inhibition show that PKA, that increases Pin1 Ser16 phosphorylation, is central to PTH secretion in parathyroid glands from rats and mice with normal renal function and from mice with SHP induced by either CKD or AKI.

To further study the effect of PKA activation on *PTH* expression, we used human embryonic kidney 293 (HEK293) cells transiently transfected with an expression plasmid carrying the human PTH gene. We have previously shown that this model recapitulates the regulation of PTH mRNA decay by calcium and calcimimetic agents through protein-*PTH* mRNA interactions (12, 53, 54). Forskolin led the expected increase in Pin1 Ser16 phosphorylation in HEK293 cells (Supplemental Fig 5) (33). Importantly, forskolin increased PTH mRNA levels at both 3 and 24 h (Fig 4E), similar to its effect to increase PTH secretion from parathyroid glands in culture (Fig 4A). Altogether, these studies show that PKA activation, that increases Pin1 Ser16 phosphorylation, mediates the increased *PTH* expression ex-vivo in parathyroid glands in culture and in transfected HEK293 cells. These findings are consistent with increased parathyroid Pin1 phosphorylation in vivo (Fig 3) and the decreased Pin1 activity (12) in kidney failure induced SHP.

### Targeting the Pin1 catalytic domain by the pharmacological inhibitor Sulfopin increases *PTH* expression in parathyroid organ cultures

To further understand the role of Pin1 in mediating *PTH* expression, we studied the effect of Pin1 inhibition by Sulfopin on PTH expression ex-vivo in parathyroid organ cultures. Sulfopin is a highly selective nanomolar-range Pin1 inhibitor that targets Cys113 in the catalytic domain (Fig 3C) (32, 37). To study the effect of Sulfopin on PTH secretion and mRNA levels, we used mice that express YFP specifically in their parathyroids (PT-*YFP*), that we have generated (Fig 5A). After fluorescence-guided microsurgery, mouse parathyroid glands with minimal adjacent thyroid tissue, were incubated in culture medium containing Sulfopin or vehicle (DMSO), as above. Pin1 inhibition by Sulfopin led to an increase in PTH secretion. Sulfopin also increased PTH mRNA levels in the micro-dissected PT-*YFP* mouse parathyroid glands, that otherwise are difficult to detect and separate form surrounding thyroid tissue (Fig 5B-C). Altogether, our data show that Pin1 pharmacologic inhibition at both the protein-binding and catalytic domains, that mimics the decreased Pin1 isomerase activity in SHP, increases PTH levels.

**Figure 5.**
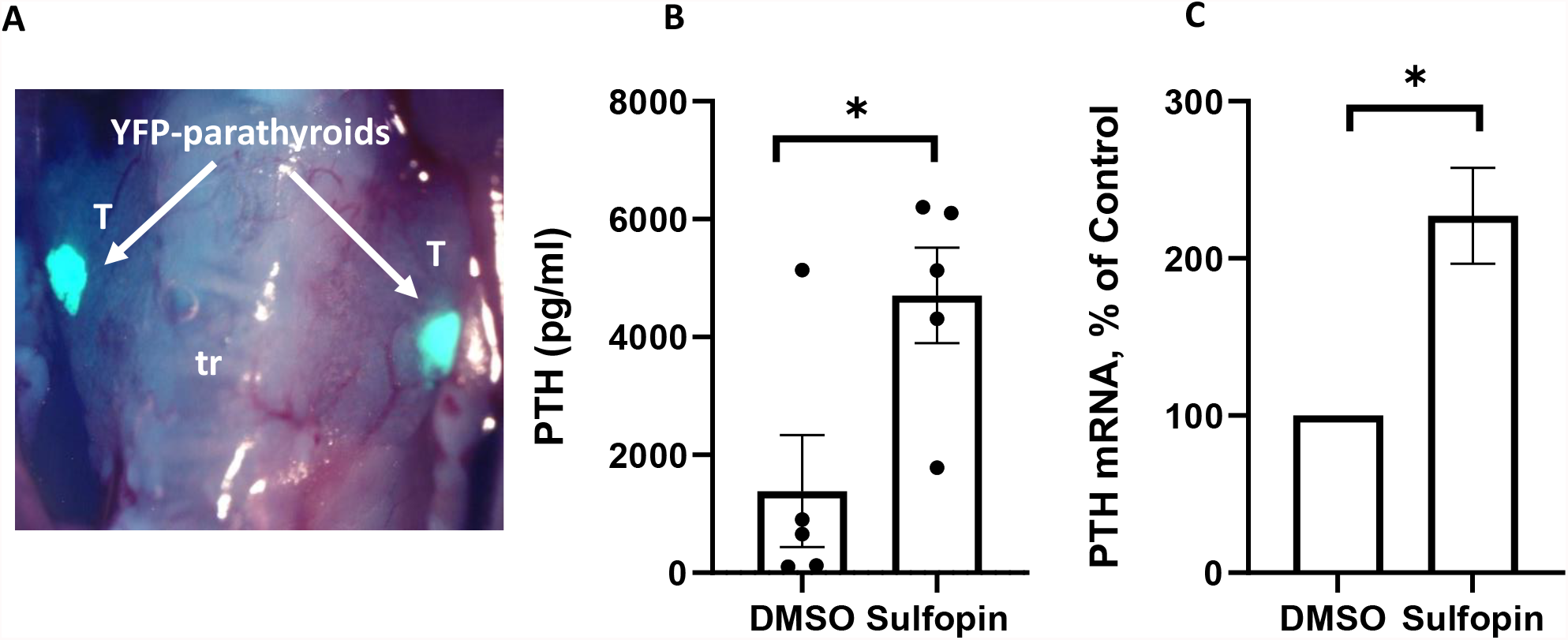
Targeting Pin1 Cys113 at the catalytic domain by the covalent Pin1 inhibitor Sulfopin increases PTH expression in mouse parathyroid glands in culture. **(A**) The exposed neck area in a mouse that shows YFP expression specifically in the parathyroids (PT-*YFP*) used for fluorescence-guided micro-surgery of the parathyroid glands with minimal contamination of the adjacent thyroid tissue. T, thyroid, Tr, trachea. (**B**) Microdissected parathyroid glands from PT-*YFP* mice were maintained in culture with Sulfopin (50 µM) or DMSO and PTH accumulated in the growth medium measured at 24 h. Results are presented as mean±SE, compared vehicle treated glands. *, p<0.05. (**C**) PTH mRNA levels normalized to β-actin were determined by qRT-PCR in pooled glands from **B** and pools form an additional repeat experiment, each composed of pooled parathyroid glands from 5 mice in each group.

### Pin1 inhibition at either the WW or the catalytic domain increases *PTH* expression through the PTH mRNA 3’-UTR 63 nt protein-binding element

We have previously identified a defined 63 nt ARE in the *PTH* mRNA 3’-UTR that is both necessary and sufficient for protein-mRNA interaction and the regulation of *PTH* mRNA stability in SHP parathyroids and in vitro when introduced in a reporter growth hormone (*GH*) expression plasmid containing the *PTH* mRNA 3’-UTR 63 nt *cis-*acting element (*GH63*) (11, 12, 53). To determine if Pin1 pharmacological inhibition by forskolin mediated PKA activation and Sulfopin increases *PTH* expression through the *PTH* mRNA ARE, we transfected HEK293 cells with a wt *GH* expression plasmid or a *GH*63 reporter plasmid (Fig 6A) (53). Forskolin increased reporter *GH63* mRNA levels, similar to its effect to increase *PTH* mRNA (Fig 4E), but had no effect on wt *GH* mRNA that did not contain the PTH mRNA element (Fig 6B). Likewise, Pin1 inhibition by Sulfopin increased *GH63* mRNA levels, but not wt *GH* mRNA levels (Fig 6C). Therefore, Pin1 inhibition induced by both PKA activation and Sulfopin increases *PTH* expression through the *PTH* mRNA 3’-UTR 63 nt protein-binding element.

**Figure 6.**
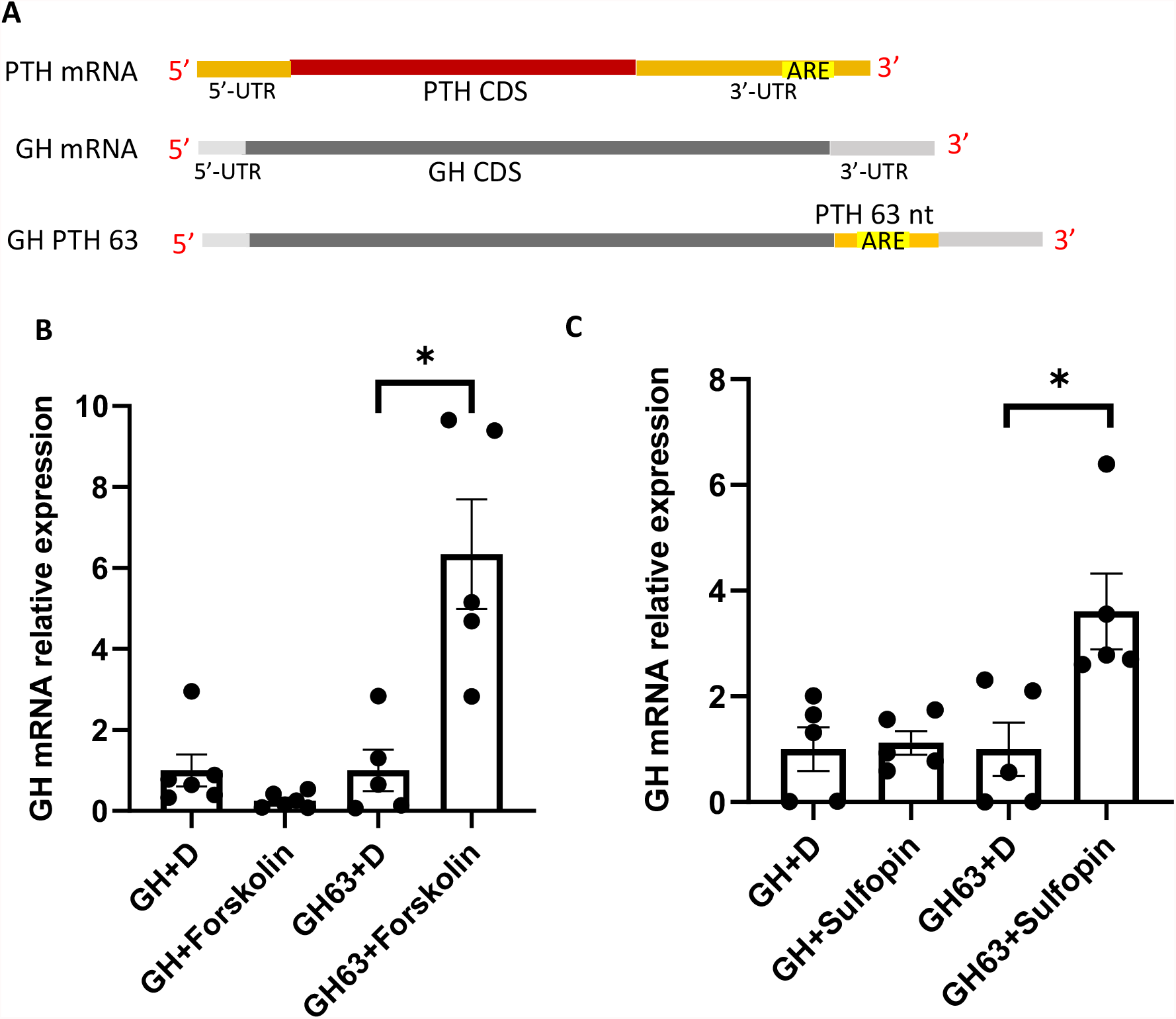
Pin1 inhibition by forskolin or Sulfopin increases PTH mRNA levels through the PTH mRNA 3’-UTR 63 nt protein-binding element. **(A**) Schematic representation of the constructs used for transfection, showing the PTH, native growth hormone (GH) and the reporter GH containing the PTH 3’-UTR AU rich 63 nt *cis* acting element (ARE) mRNAs (GH63). CDS, coding sequence; UTR, un-translated region. (**B-C**) HEK293 cells were transiently transfected with expression plasmids for GH or GH63 (GH63). Cells were treated with Forskolin (5 µM) (**B**) or Sulfopin (5 µM) (**C**) or vehicle (DMSO, +D). PTH mRNA levels were measures at 24 h by qRT-PCR, normalized to TATA box binding protein (TBP). Results are presented as mean±SE of fold change, compared to GH mRNA in vehicle treated cells. *, p<0.05 inhibitor vs vehicle.

### Pin1 inhibition by both forskolin and Sulfopin alters over-all protein-*PTH* mRNA 3’-UTR interaction

To characterize the PTH mRNA 3’-UTR binding proteins and the effect of Pin1 inhibition on protein-RNA binding, we performed liquid chromatography-mass spectrometry (LC-MS) analysis of *PTH* mRNA 3’-UTR interacting proteins from HEK293 cells incubated with forskolin, Sulfopin or vehicle. Proteins were pulled-down using biotinylated in vitro-transcribed human *PTH* mRNA 3’-UTR immobilized onto streptavidin beads and transcript-bound proteins were eluted and across all extracts, MS identified 610 proteins (post filtering of common contaminants), of which 82 proteins (13.4%) were detected at levels significantly above the background recovered from uncoated streptavidin beads (Supplemental Fig 6A). The majority of these (53%) were RNA binding proteins and many others were involved in mRNA regulation and processing (Fig 7A, and Supplemental Fig 6B). The top enriched RNA binding proteins were PRKRA (an RNA-dependent protein kinase activator) and IGF2BP1 (a well-studied mRNA-BP, which inhibits both mRNA translation and degradation, depending on its binding target). Additional highly enriched proteins were KSRP and AUF1 (HNRNPD), the two PTH mRNA-interacting proteins central for increased *PTH* expression in SHP (11, 14), confirming the validity of the assay. Of interest, Pin1 was also pulled-down specifically by the PTH mRNA 3’-UTR (Supplemental Fig 6A). Enrichment of several pulled-down proteins was boosted in extracts recovered from cells incubated with either of the Pin1 inhibitors, and the effect of inhibitors was largely similar (Fig 7B and Supplemental Fig 6C-D). The most significantly amplified proteins were APEH, NEU2, GLOD4, NAP1l4, ILF2/3, PRKRA, IGF2BP1/3 and STAU2. In the list of pulled-down proteins (82 significantly above background and 73 with borderline significance, Supplemental Fig 6A), those proteins further intensified in the presence of Pin1 inhibitors (n=31) were enriched with the following annotations: double-stranded RNA binding, protein binding, poly(A) RNA binding and intracellular ribonucleoprotein complex. Thus, Pin1 inhibition results in altered protein-*PTH* mRNA interactions. The roles of these PTH-interacting proteins in regulation of *PTH* expression in the context of Pin1 inhibition in vitro and in parathyroids in SHP remains to be determined.

**Figure 7.**
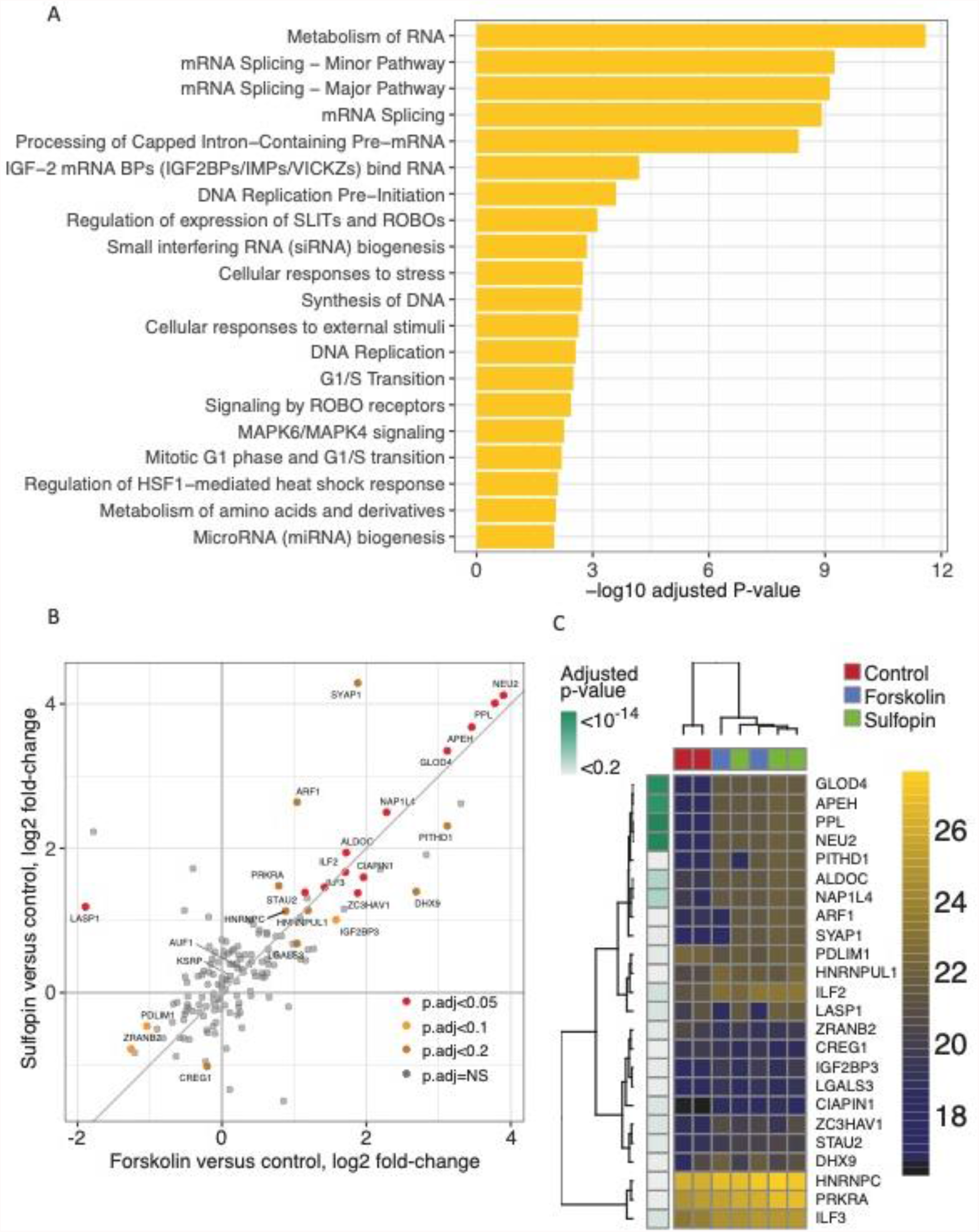
Mass-spectrometry based discovery of PTH mRNA binding proteins and the response to Pin1 inhibition mimicking experimental CKD. (**A**) Characteristics of the PTH 3’-UTR specifically bound (pulled-down) proteins when evaluated for enrichment in Reactome database pathways at an adjusted p-value cutoff 0.01. Fig S5B expands this analysis to additional data sources. (**B**) Results of differential pulldown analysis from Pin1-inhibited cell extracts versus control cells, wherein the y-axis shows the effect of Sulfopin and the x-axis shows the effect of forskolin. (**C**) Heat map showing intensity values of the same set of differentially-pulled-down proteins.

### PKA activated Pin1 inhibition decreases Pin1-KSRP interaction

To understand the effect of Pin1 inhibition on the *PTH* mRNA decay promoting protein KSRP, we first studied Pin1-KSRP protein-protein interaction after Pin1 inhibition. Cells were transiently transfected with *FLAG-KSRP* expression plasmid and then incubated with forskolin, Sulfopin or vehicle. At 24 h extracts were prepared and immunoprecipitated with Pin1 antibody. Western blots showed that FLAG-KSRP was pulled-down by Pin1, as expected (Supplemental Fig 7) (12). Forskolin decreased Pin1-FLAG-KSRP interaction, consistent with forskolin induced Pin1 Ser16 phosphorylation that targets the Pin1 protein-binding domain. Sulfopin acting on the catalytic domain of Pin1, had no effect on Pin1-KSRP interaction (Supplemental Fig 7). Therefore, PKA activation leads to dissociation of Pin1 from KSRP in HEK293 cells.

### Pin1 induced changes in KSRP phosphorylation alter *PTH* expression

Phosphoproteome analysis showed that KSRP is hyperphosphorylated at Ser181, and Thr101 in parathyroids fromkidney failure rats (Fig 2C). We have previously shown that KSRP is a target protein for Pin1 and that Pin1 over-expression leads to KSRP Ser181 dephosphorylation that promotes *PTH* mRNA decay (12). KSRP Thr692 phosphorylation impairs KSRP’s decay-promoting activity on a variety of target mRNAs (55). Ser181, Thr101 and Thr692 all precede Pro as expected from target sites for Pin1 isomerase activity (Fig 2C and not shown). To characterize the impact of these KSRP pSer/Thr-Pro phosphorylation sites on *PTH* mRNA levels, HEK293 cells were co-transfected with expression plasmids for *PTH* and wt *KHSRP* or *KHSRP* phosphonull mutants. Co-transfection of *PTH* and wt *KSRP* expression plasmids led to the expected decrease in *PTH* mRNA (Fig 8B and D), consistent with the *PTH* mRNA decay promoting activity of KSRP (12). Interestingly, Ser181Ala, Thr100Ala and Thr692Ala single phosphorylation null mutants had a similar effect as wt KSRP to decrease *PTH* mRNA, and forskolin only partially reversed this effect with expression of the KSRP constructs (Fig 8B&C). In contrast to the KSRP single phosphorylation null mutants, the ability of wt KSRP to decrease *PTH* mRNA was magnified by over-expression of KSRP Ser181Ala;Thr100Ala double mutant and more so by the Ser181Ala;Thr100Ala;Thr692Ala triple mutant (Fig 8D). Therefore, KSRP dephosphorylation at more than one Ser/Thr-Pro motif is required to maximize the KSRP mediated decrease in *PTH* expression. Forskolin activated PKA induced Pin1 phosphorylation, which would result in KSRP phosphorylation (12), did not restore the increase in *PTH* expression when the double or triple KSRP phosphorylation null mutants were over-expressed (Fig 8D). Therefore, the increase in *PTH* expression by Pin1 phosphorylation and inhibition is dependent on KSRP phosphorylation.

**Figure 8.**
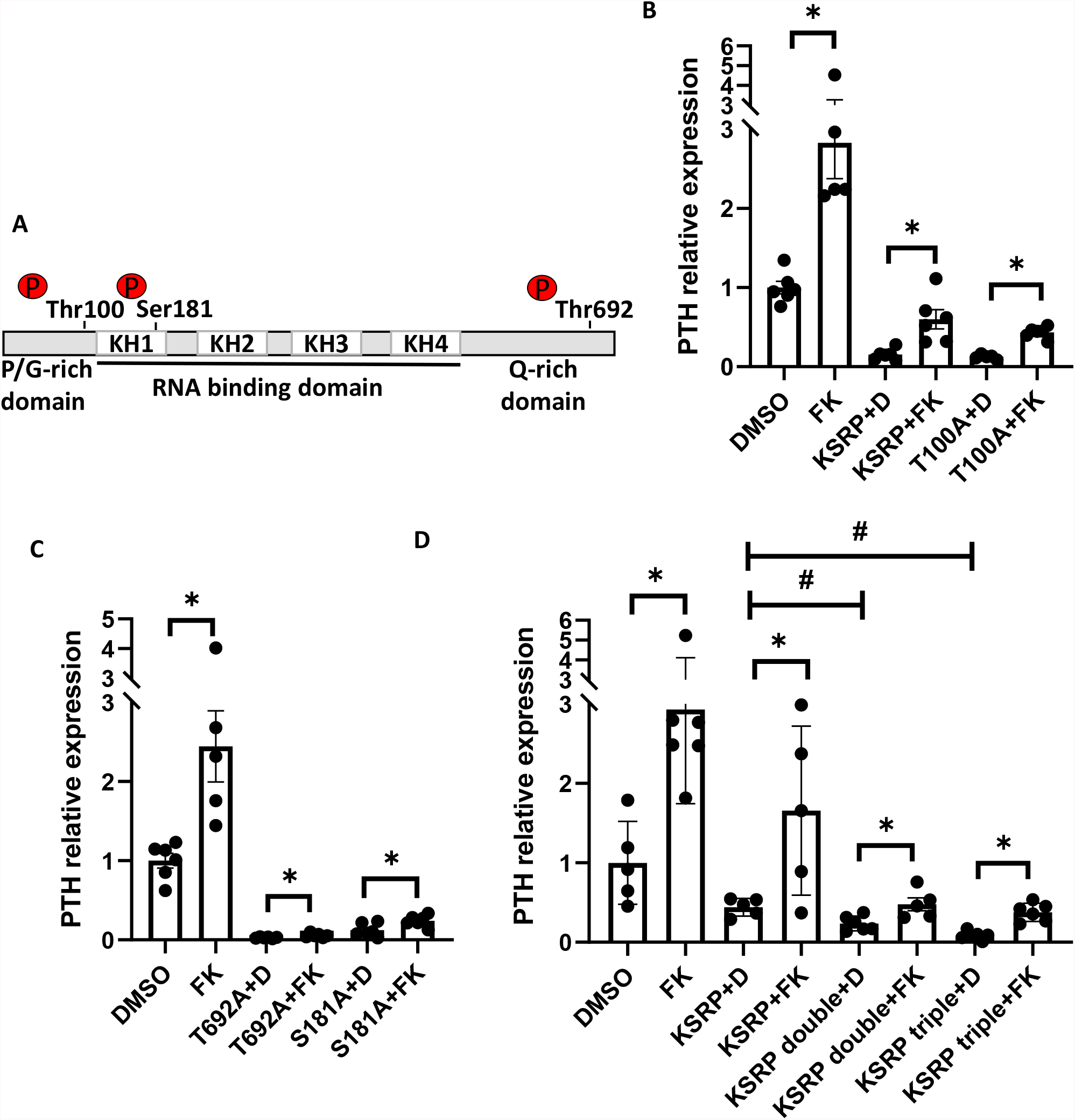
PKA activation increases PTH expression through KSRP phosphorylation. A. Schematic representation of the primary structure of KSRP, including the four hnRNP K homology (KH) domains and position of the phosphorylation sites relevant to this study. (**B-D**) HEK293 cells were transiently transfected with expression plasmids for human *PTH* with empty vector or native *KSRP* or *KSRP* phosphorylation null single mutants Thr100Ala (T100A) (**B**); Thr692Ala (T692A) or Ser181Ala S181A) (**C**) or *KSRP*, and *KSRP*Ser181Ala;Thr100Ala (double) or Ser181Ala;Thr100Ala;Thr692Ala (triple) mutants **(D**). Transfected cells were incubated with forskolin (FK) or vehicle DMSO (+D). PTH mRNA levels at 24 h, normalized to GAPDH, were determined by qRT-PCR. Results are presented as mean±SE of fold change, compared to cells transfected with PTH expression plasmid and a control empty vector. *, p<0.05 forskolin vs DMSO; #, vs KSRP+D.

We next asked whether Pin1 inhibition by Sulfopin and KSRP phosphorylation act via the *GH-PTH63* (*GH63*) mRNA element. We therefore co-transfected cells with expression plasmids for GH63 and either KSRP or the KSRP Ser181Ala;Thr100Ala;Thr692Ala triple phosphorylation null mutant. Sulfopin increased *GH63* mRNA levels in wt KSRP overexpressing cells (Supplemental Fig 8), similar to its effect without KSRP over-expression (Fig 6C). Importantly, Pin1 inhibition failed to increase *GH63* mRNA levels in cells over expressing KSRP triple phosphorylation null mutant (Supplemental Fig 8). Therefore, Pin1 inhibition by Sulfopin increases PTH expression in a manner that is also dependent on KSRP phosphorylation, at least in 2 phosphorylation sites. The Pin1-KSRP axis and the *PTH* mRNA 3’-UTR *cis* element are central to this increase in PTH expression. These findings are consistent with loss of Pin1 activity in SHP parathyroids that increases PTH expression through modified *PTH* mRNA-KSRP interaction (12).

## Discussion

Secondary hyperparathyroidism is a frequent complication and a key player in the devastating clinical consequences of CKD. The study of the molecular signals that induce SHP in CKD has been hampered by the lack of an appropriate cell culture system and the minute size of the parathyroid glands in rodent models. Here we aim to elucidate the signaling pathways and underlying molecular mechanisms involved in the stimulated PTH synthesis and secretion in SHP using in vivo kidney failure models combined with ex-vivo and cell culture assays. Mass spectrometry analysis identified changes in parathyroid proteome and phosphoproteome profiles induced by experimental CKD. To the best of our knowledge, this is the first attempt to characterize the global changes in protein expression and phosphorylation induced by kidney failure in the minute rat parathyroid glands. We used the experimental model of CKD where kidney failure is induced by an adenine high phosphorus diet. This model allows to study of the effects of mild renal failure combined with hyperphosphatemia induced SHP, already at 2 weeks of the diet. At this time point there is no significant decrease in parathyroid VDR, CaR and klotho protein levels, as can be found later, at 6 weeks of the adenine high phosphorus diet (51). Indeed, our MS analysis did not detect changes in the levels of these proteins that participate in the establishment of SHP. Nevertheless, our data highlight new potential pathways that may be involved in the early CKD induced changes in parathyroid function resulting in SHP.

Phosphoproteome analysis identified up- and down-regulated phosphosites, with more phosphosites upregulated than downregulated, suggesting an increase in global protein phosphorylation. Increased intracellular phosphate concentrations were reported in uremic SHP (43). We show enrichment of mTOR signaling-related proteins. Indeed, mTOR activation is central to the development of SHP and parathyroid cell proliferation induced by either kidney failure or hypocalcemia in rats, confirming the validity of the assay (40). Kinase activity scores that sum changes in phosphorylation of target proteins, pointed to increased activity of several kinases in experimental CKD parathyroids. Among them, PKA increased activity was demonstrated. PKA inactivates Pin1 isomerase activity by targeting Pin1 Ser16 phosphorylation in the WW domain, leading to dissociation of Pin1 from its substrates (33). In the context of PTH regulating pathways, PKA activation could mediate the decreased parathyroid Pin1 activity that occurs in SHP induced by either kidney failure or prolonged hypocalcemia (12). We then looked at global differences between parathyroid proteins from adenine fed kidney failure rats, with enriched and depleted phosphosites (**Fig 2A**). Indeed, the change in potential phosphorylated Ser/Thr-Pro that are Pin1 target sites, in CKD rat parathyroid extracts, highlights a role for Pin1 in SHP.

Among the proteins identified by the phosphoproteome, KSRP was phosphorylated in kidney failure rat parathyroids at potential targets for Pin1 isomerize activity. We have previously shown that KSRP interacts with and is a target protein for Pin1. Pin1 over-expression in transfected human cells decreases KSRP phosphorylation at Ser181 which resulted in decreased *PTH* mRNA stability and levels (12). The decreased Pin1 activity in CKD parathyroids (12) could increase KSRP phosphorylation. Indeed, kidney failure led to parathyroid KSRP Ser182 (Ser181 in human KSRP) hyperphosphorylation by phosphoproteome analysis. In addition to Ser181, Pin1 phosphorylation at Thr101, was also increased in CKD parathyroids. KSRP Ser481 phosphorylation, another yet unidentified potential Pin1 target, was decreased. These KSRP post-translational modifications may be due to the decreased Pin1 activity in SHP parathyroids and could alter KSRP-*PTH* mRNA interaction and contribute to the increased *PTH* mRNA levels in SHP.

Of interest, Ser181 KSRP and Pin1 phosphorylation was also identified in PTH-stimulated opossum kidney (OK) proximal tubule cell culture model, by MS phosphoproteome characterization (20). It is intriguing that KSRP and Pin1 are involved in both the kidney and parathyroid, two organs central to mineral metabolism. Recently, mRNA profiles of porcine parathyroid glands were performed in a long-term dietary phosphorus intervention, by keeping pig offspring on distinct mineral phosphorus levels throughout fetal and postnatal life. RNA sequencing data and resulting molecular pathways of parathyroid glands showed that PTH abundance is controlled via Pin1, CaSR, MAfB, PLC and PKA signaling to regulate *PTH* expression, stability, and secretion. Parathyroid glands revealed lowered *Pin1* mRNA abundance in animals fed a low phosphorus diet with no change in the expression of *KSRP* by post-weaning diets (56).

Based on phosphoproteome analysis, that showed PKA activation and changes in KSRP phosphorylation in kidney failure parathyroids, and our previous studies on KSRP and Pin1 interaction in the parathyroid, we chose to study the parathyroid KSRP-Pin1 axis and its effects on PTH expression in CKD SHP. We show that both experimental CKD and AKI lead to increased parathyroid Pin1 phosphorylation at Ser16 and 71. This specific hyperphosphorylation is consistent with the decreased Pin1 isomerase activity in other systems (34, 35) and with the decreased Pin1 activity in parathyroids of rats with hypocalcemia or CKD induced SHP (12). PKA activation in SHP parathyroids would contribute to this Pin1 Ser16 phosphorylation and decreased isomerase activity. We then studied the mechanisms of decreased Pin1 activity both ex-vivo in parathyroid glands in culture and in transfected cells, as there is no functional parathyroid cell line. Pharmacologic inhibition of Pin1 by forskolin induced PKA activation leads to Ser16 Pin1 phosphorylation at the protein-binding WW domain (33). Indeed, forskolin increased *PTH* expression in both parathyroid organ cultures and transfected cells. PKA inhibition, that would leave Pin1 unphosphorylated and active, had the opposite effect to decrease *PTH* mRNA levels. Forskolin has been shown to increase cellular cAMP content and parathyroid hormone release in dispersed primary bovine parathyroid cells (57). Cholera toxin which activates adenylate cyclase and indirectly stimulates PKA, increased *PTH* mRNA and PTH secretion in these cells (58). The mechanism of increased *PTH* expression by PKA activation was not studied at the time, but may be due to the activated PKA induced Pin1 Ser16 phosphorylation and decreased Pin1 activity, as suggested by our current findings.

Inhibition of Pin1 by targeting C113 in the catalytic domain with Sulfopin had similar effects as PKA activation, to increase *PTH* expression. Therefore, pharmacologic inhibition, at both the WW and the catalytic domain, that mimics the decreased Pin1 activity in vivo in the CKD SHP parathyroids, leads to increased *PTH* expression also when added directly in vitro. These effects were both dependent upon the *PTH* mRNA 3’-UTR *cis* acting element. The effect of Pin1 inhibition to increase PTH expression is consistent with our previous findings that *Pin1*^*-/-*^ mice have increased parathyroid gland PTH mRNA and protein levels and circulating serum PTH concentrations (12). Altogether these findings support a role for Pin1 inhibition to increased PTH expression. To better understand the effect of Pin1 inhibition on *PTH* expression and mRNA stability, we identified by Mass spectrometry the *PTH* mRNA 3’-UTR interacting proteins and the effect of Pin1 inhibition on mRNA binding. We identified many proteins that are involved in mRNA fate, among them KSRP, AUF1 and Pin1.

KSRP over-expression that induces PTH mRNA decay and decreases PTH expression in transfected cells (12) was more potent in decreasing *PTH* mRNA levels when KSRP phosphorylation null mutations at Pin1 potential target site were introduced. Moreover, the effect of Pin1 pharmacologic inhibition to increase *PTH* expression was dependent upon KSRP phosphorylation at Ser181 and Thr100 and more so when an additional phosphorylation site Thr692 is present. p38 MAPK-mediated KSRP phosphorylation at Thr692 Pin1 potential site (pThr692-Pro693) impairs KSRP-RNA interactions and increases target mRNA abundance in a variety of target mRNA, demonstrating that KSRP phosphorylation is crucial for its mRNA decay promoting capabilities (26). These findings are consistent with the decreased Pin1 activity in vivo in SHP that correlates with increased *PTH* expression (12) and the increased KSRP phosphorylation at Ser181 and Thr100 shown in the phosphoproteome analysis. Of interest, in a human study of single nucleotide polymorphisms (SNP) in the Pin1 gene promoter, in the Chinese Han population in Northwest China, Pin1 C667T genetic variants were associated with CKD SHP (59).

In summary, we show that experimental CKD induces global changes in expression and phosphorylation of parathyroid proteins. Specifically, the PKA pathway is activated and KSRP is hyperphosphorylated in kidney failure parathyroids. Kidney failure also induces Pin1 phosphorylation at residues that inhibit its isomerase activity, explaining the decreased Pin1 activity we have previously shown in SHP parathyroids (12). The decreased Pin1 activity and the resulting KSRP phosphorylation at Pin1 target sites would alter protein-*PTH*-mRNA interactions and hence increase PTH expression. These effects can be mimicked in vitro in parathyroid glands in culture and in transfected human cells by pharmacological phosphorylation and inhibition of Pin1. We propose that impaired renal function leads to parathyroid Pin1 phosphorylation and decreased Pin1 activity. This increases PTH expression in SHP in a manner that is dependent on KSRP phosphorylation at Pin1 target residues and the PTH mRNA *cis* acting element (Fig 9). Pin1 phosphorylaiton and the resulting loss of Pin1 activity in SHP parathyroids triggers the parathyroids to increase PTH expression through modified Pin1 and *PTH* mRNA-KSRP interaction and KSRP phosphorylation.

**Figure 9.**
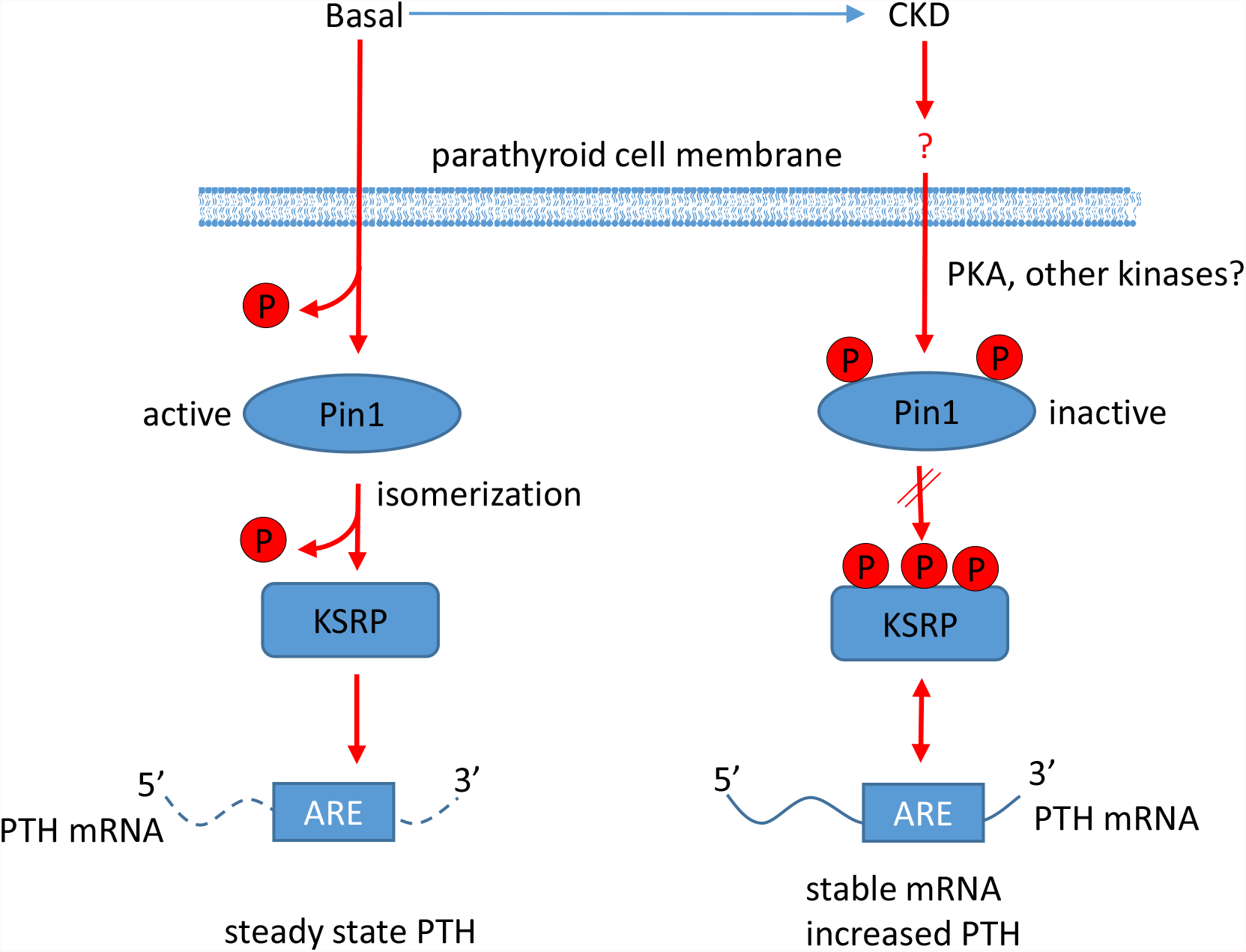
Model for the changes that renal failure induces in the parathyroid cell that lead to increased PTH gene expression in SHP. Under basal conditions, Pin1 is unphosphorylated at Ser16 and 71 and thus active, leading to conformational change leading to decreased KSRP phosphorylation. Unphosphorylated KSRP binds to the adenine uridine rich element (ARE) in the PTH mRNA 3’-untranslated region and along with other PTH mRNA binding proteins determines steady state PTH levels. CKD induces parathyroid Pin1 phosphorylation and hence decreased isomerase activity, leaving KSRP phosphorylated at three Pin1 target sites. Phosphorylated KSRP fails to bind PTH mRNA and induce mRNA decay, resulting in increased PTH mRNA stability and high serum PTH levels.

## Materials and Methods

### Animals, housing and diets

Experimental chronic kidney disease (CKD) was induced in male Sprague-Dawley rats (150–170 g) by an adenine-rich (0.75%) high-phosphorus (1.5%) diet (ENVIGO, Harlan Laboratories, Madison, WI) given for 14 d (12). Controls received regular chow containing 0.7% phosphorus. CKD in mice was induced in male C57BL/6 mice at 11–12 wk of age by a moderate (0.3%) adenine-rich high-phosphorus (1.2%) diet (ENVIGO) given for 14 d (52, 60). Acute kidney injury (AKI) was induced in male C57BL/6 mice at 11–12 wk of age by a single injection of folic acid (240 mg/kg in vehicle, 0.15 mol/l NaHCO3, pH 7.4, Sigma-Aldrich, St. Louis, MO) (9, 10). Control mice were injected with vehicle. Mice were analyzed at 24 and 48 h. In some experiments, PT-*EYFP* (parathyroid specific enhanced Yellow Fluorescent Protein) mice were used to allow identification and micro-dissection of the mouse parathyroid glands for organ culture experiments. We generated the PT-*EYFP* mice by Cre-Lox recombination, mating PT-*Cre* mice where the Cre recombinase is driven by the human PTH promoter [FVBTg(PT-Cre); Jackson Laboratory, Bar Harbor, ME] (2) with *R26-stop-EYFP/R26R-<u>EYFP</u>*, mice where a loxP-flanked STOP sequence followed by the *EYFP* was inserted into the Gt(ROSA)26Sor locus (61). Total DNA was extracted from tail samples of offspring and genotyping was performed by PCR. The primers used were, for Cre: Fw: 5’-TGCCACGACCAAGTGACAGC-3’and Rev: 5’-CCAGGTTACGGATATAGTTCATG-3’ (52) and for EYFP: ROSA 26R 5’-AAAGTC GCTCTGAGTTGTTAT-3’; BTG 60: 5’-GAAAGACCGCGAAGAGTT TG-3’ and BTG 62: 5’-TAAGCCTGCCCAGAAGACTC-3’. All animals had free access to food and drinking water. Experiments were approved by the Institutional Animal Care and Use Committee of the Hebrew University-Hadassah medical school (authorization numbers MD-18-15408, 18-15610).

### Parathyroid glands in organ culture

The 2 glands from micro-dissected mouse thyro-parathyroid tissue, parathyroid glands from PT-*EYFP* mice or rat parathyroids (n=6) were maintained in 2 ml Eppendorf tubes with needle-punctured caps for aeration, containing 1 ml DMEM (Gibco Life Technologies, Carlsbad, CA) supplemented with 10% fetal bovine serum, L-glutamine, and penicillin-streptomycin (Gibco Life Technologies). Forskolin at a concentration of 100 µM, H89 (N-[2-p-bromocinnamylamino-ethyl]-5-isoquinolinesulphonamide) at 150 µM, Sulfopin at 50 µM or vehicle (DMSO) were added to the medium. The tubes were placed in a CO_2_ incubator with constant rocking (52, 62). Medium (100 µl) was collected at the time points indicated and analyzed for PTH. In some experiments, thyroid-parathyroid glands were removed from adenine high–phosphorus diet induced CKD, AKI and control mice.

### Serum biochemistry and PTH levels

Serum was analyzed for calcium and blood urea nitrogen (BUN) using QuantiChrom kits (BioAssay Systems, Hayward, CA). Serum phosphate was analyzed using a Stanbio Phosphorus Liqui-UV kit (Stanbio Laboratories, Boerne, TX). Serum PTH or PTH secreted to growth medium in organ cultures was measured using rat or mouse 1–84 Intact PTH ELISA kit (Quidel, Athens, Ohio).

### RNA-protein extraction

Thyro-parathyroid glands in organ culture were removed from the medium at the end of the incubation period and homogenized using a bead-beater. RNA was extracted from pools of glands form 6 mice using TRIzol Reagent (Invitrogen, Carlsbad, CA). RNA from HEK293 cells was also extracted with TRIzol Reagent (12). Protein extracts for immuno-precipitation (IP) in cultured HEK293 cells were prepared using RIPA buffer containing 150 mM NaCl, 1% NP40, 0.5% sodium deoxycholate, 0.1% SDS and protease inhibitors cocktail (Roche, Mannheim, Germany). Antibodies used were against Pin1 (Abcam, Cambridge MA) and FLAG, (Sigma-Aldrich).

### Mass Spectrometry (MS) for proteomics and phosphoproteomics was performed at the Technion Israel Institute of Technology, Haifa, Israel

For proteome analysis, pooled parathyroid tissue samples from 2 rats in every group in triplicate were homogenized in lysis buffer and sonicated. The proteins were digested in 1 M urea with modified trypsin (Promega, Madison, WI) at a 1:50 enzyme-to-substrate ratio, overnight at 37° C. The tryptic peptides were desalted and re-suspended in 40% acetonitrile (ACN), 6% TFA (Trifluoroacetic acid), and enriched for phosphopeptides on titanium dioxide beads. Bound peptides were eluted with 20% ACN with 325 mM Ammonium Hydroxide followed by 80% ACN with 325 mM Ammonium Hydroxide. The resulting peptides were desalted and analyzed by LC-MS/MS on a Q-Exactive plus mass spectrometer (Thermo-Fisher scientific, Pittsburgh, PA) fitted with a capillary HPLC (easy nLC 1000, Thermo-Fisher scientific). Mass spectrometry was performed in a positive ion mode (at mass range of m/z 350–1800 AMU and resolution 70,000) using repetitively full MS scan followed by collision induces dissociation (HCD, at 35 normalized collision energy) of the 10 most dominant ions (>1 charges) selected from the first full MS scan. For the phosphoproteomic profiling, samples 2 pools of 6 rats from control and kidney failure rats underwent phosphoenrichment using Immobilized Metal (Fe^+3^) Affinity Chromatography on a robotic system (Bravo) prior to LC-MS/MS analysis.

### Mass spectrometry data analyses

The mass spectrometry data was analyzed using the MaxQuant software 1.5.2.8 (www.maxquant.org, (63)) for peak picking identification and quantitation using the Andromeda search engine, searching against the rat proteome from the Uniprot database with mass tolerance of 20 ppm for the precursor masses and 20 ppm for the fragment ions. Methionine oxidation, phosphorylation (STY) and protein N-terminus acetylation were accepted as variable modifications and carbamidomethyl on cysteine was accepted as static modifications. Minimal peptide length was set to six amino acids and a maximum of two miscleavage events was allowed. Peptide- and protein-level false discovery rates (FDRs) were filtered to 1% using the target-decoy strategy. The protein table was filtered to eliminate the identifications from the reverse database, and common contaminants. The data was quantified by label free analysis using the MaxQuant software, based on extracted ion currents (XICs) of peptides enabling quantitation from each LC-MS run for each peptide identified in any of experiments.

The rat parathyroid MS proteomics and phosphoproteomics data have been deposited to the ProteomeXchange Consortium via the PRIDE partner repository with the dataset identifiers PXD029368 and PXD029401, respectively (64). For the primary statistical analysis of the MaxQuant *proteomic* data we uploaded proteinGroups.txt and evidence.txt files to the ProteoSign webserver (65). Supplemental Table S1 and Fig 1 present ProteoSign’s differential protein expression results and Supplemental Fig 1 shows ProteoSign’s enrichment results, processed with g:Profiler (66) and plotted using ggplot2 (https://ggplot2.tidyverse.org) for R. MaxQuant-identified *phosphoproteins*, with the detectable proteome as background, subjected to enrichment analysis using DAVID (67), and the results displayed with ggplot2. A recently described R package, PhosR, was applied for differential phosphorylation analysis, for phosphosite- and gene-centered enrichment analyses of differentially phosphorylated sites and for kinase perturbation analysis (42). Finally, we searched for significant sequence motif alterations in phosphosites between Adenine and Control with Two Sample Logo approach, using a webserver interface (48).

### Cell cultures and transient transfection

HEK293 cells were transiently transfected in 24-well plates for RNA analysis and in 10 cm plates for protein extractions and IP respectively using a TransFectinTM reagent (Bio-Rad, Hercules, California). 4 h after transfection, the cells from the 24 well plates were trypsinized and re-seeded in 96 well plates. The next day cells were treated with vehicle (DMSO), forskolin or Sulfopin at 5 µM, for the indicated time points. RNA was extracted with TRIzol (Invitrogen) for qRT-PCR analysis.

### Plasmids

The human PTH gene, including exons and introns in pcDNA3 was previously described (54). pCMV-Tag 2B plasmid containing FLAG-full length KHSRP or the KHSRP phosphorylation null mutants Thr692Ala, Thr100Ala where kindly provided by R. Gherzi (IRCCS Ospedale Policlinico San Martino, Genova, Italy). Ser181Ala, Ser181Ala;Thr100Ala double mutant and Ser181Ala;Thr100Ala;Thr692Ala triple mutant were prepared using the QuikChange Lightning Site-Directed Mutagenesis Kit (Agilent Technologies, Santa Clara, CA, US) according to the manufacturer’s instructions. The GH expression plasmid was kindly provided by O. Meyuhas (Hebrew University-Hadassah Medical School, Jerusalem, Israel). The *GH-PT*H mRNA 63nt plasmid (*GH63*) containing the 63nt rat *PTH* mRNA ARE cloned between the 3′ of the *GH* mRNA coding sequence and the *GH* mRNA 3′ UTR was previously described (11, 53). Empty control vector pcDNA3 (Invitrogen) was used as control. GST-Pin1 expression plasmid was a kind gift from K.P. Lu (Beth Israel Deaconess Medical Center, Harvard Medical School, Boston, MA).

### Quantitative RT-PCR analysis

cDNA was synthesized using a qScript cDNA Synthesis Kit (Quanta bio, Beverly, MA) and quantitative (q) PCR analysis performed with PerfeCTa SYBR Green FastMix, Low ROX (Quanta bio) in a ViiA 7 Fast Real-Time PCR System (Applied Biosystems Waltham MS). Primers were as follows: mouse PTH: Fw: 5’-ATCCCTTTGAGAGTCATTG-3’ and Rev: 5’-GTTTGGGTAAGAAGACAGAC-3’; mouse β-actin: Fw: 5’-CCTAGGCACCAGGGTGTGAT-3’ and Rev: 5’-TCAGGGTCAGGATACCTCTCTTG-3’; human PTH: Fw: 5’-CAGATTTCCCATCCGATTTT-3’ and Rev:5’-GGGTCTGCAGTCCAATTCAT-3’; human GAPDH: Fw: 5’-TCGGAGTCAACGGATTTG-3’ and Rev: 5’-CAACAATATCCACTTTACCAGAG-3’; human TATA-Box Binding Protein (TBP): Fw:5’-CACTCACAGACTCTCACAAC-3’ and Rev: 5’-CTGCGGTACAATCCCAGAACT-3’; human GH: Fw: 5’-CTTATCCAGGCTTTTTGAC-3’ and Rev: 5’-TTAAACTCCTGGTAGGTGTC-3’.

### Pin1 phosphorylation assay

GST-tagged bait proteins were purified using Pierce GST Protein Interaction Pull-Down kit (Thermo-Fisher Scientific). To examine recombinant Pin1 phosphorylation, parathyroids from control and CKD rats were micro-dissected and homogenized using a bead beater (a pool from 11-13 rats in each group) in 80 mM β-glycerophosphate, 20 mM EGTA, 15 mM MgCl_2_, 50 mM NaVO_4_, and a protease inhibitor cocktail (Roche). The lysates were centrifuged for 5 min at 4° C and the kinase assay performed at 37° C for 30 min in a 50 μl reaction volume containing 50 mM Tris-HCl (pH 7.5), 10 mM MgCl_2_, 50 mM KCl, 1 mM DTT, 1 mM EGTA, 0.16 mCi/ml [γ-^32^P]ATP, parathyroid extracts (10-20 µg) and GST or GST-Pin1 bound beads (6 µg). The beads were then washed and the proteins eluted with reduced glutathione (10 mM). Eluted proteins were run on SDS-PAGE. ^32^P-labeled GST-Pin1 was detected by autoradiography and GST/GST-Pin1 by Coomassie blue staining of the gels (28). Protein bands were quantified with Quantity One (Bio-Rad).

### Immunofluorescence staining and quantification

Micro-dissected rat parathyroid glands and mouse thyro-parathyroid tissue were embedded in paraffin and sections prepared. In some experiments, HEK293 cells were grown in 24 well plates. The cells were treated with 5 µM forskolin for 1 h and then fixed with methanol for 5 min. Immunostaining of glands or cells was performed using the following primary antibodies diluted in Cas block (Zymed Laboratories, San Francisco, CA): p-Pin1 Ser71 (1:1000, a kind gift from KP Lu, Harvard Boston) (34), p-Pin1 Ser16 (1:300, BioSS, Woburn, MA), total Pin1 (1:500 Cell Signaling, Beverly, MA), PTH (1:1000, Bio-Rad, Hercules, California). Fluorochrome–conjugated secondary antibodies Cy3 and Cy5 (Bethyl, Montgomery, TX) and nuclear staining SYTOX (Life Technologies) were used for detection. Images were obtained using a Fluoview 1000 Olympus florescence microscope. Immunofluorescence staining was quantified using ImageJ (NIH, Bethesda, Maryland). Immunofluorescence staining of HEK293 cells was visualized using the Spinning disk confocal microscopy (SDCM) which improves the speed of image acquisition and NIS elements software (Nikon, Melville, NY) was used for quantification.

### Biotinylated RNA protein pull-down assay for Mass Spectrometry was performed at the Weizmann Institute of Science, Rehovot

In vitro transcribed RNA was prepared from linearized plasmid containing the human *PTH* 3’-UTR (68) using a Biotin RNA Labeling Mix (Roche) and T7 RNA polymerase. Biotinylated RNA protein pull-down was performed according to the protocol of Panda et al (69). Biotinylated RNA was incubated with extracts (1 mg) from HEK293 cells treated with forskolin, Sulfopin or vehicle DMSO, as described above, in triplicate. Non-biotinylated RNA was transcribed and incubated with cell extracts as background control. Protein-RNA complexes were pulled down using streptavidin-coated beads and eluted with PEB buffer at 100° C for 10 min. The samples were analyzed by liquid chromatography-mass spectrometry (LC-MS). The samples were subjected to tryptic digestion using an S-trap. The resulting peptides were analyzed using Waters HSS-T3 column on nanoflow liquid chromatography (nanoAcquity) coupled to high resolution, high mass accuracy mass spectrometry (Q Exactive Plus). Each sample was analyzed on the instrument separately in a random order in discovery mode. Raw data was processed with MaxQuant v1.6.6.0. The data was searched with the Andromeda search engine against the human proteome database appended with common lab protein contaminants and the following modifications: Fixed modification-cysteine carbamidomethylation. Variable modifications-methionine oxidation, protein N-terminal acetylation, asparagine- or glutamine-deamidation, and serine-, threonine or tyrosine-phosphorylation. These MS proteomics data have also been deposited to the ProteomeXchange Consortium with the dataset identifier PXD029456. Data filtration, transformation, normalization, missing value imputation and subsequent differential expression analysis were performed using DEP package (Differential Enrichment analysis of Proteomics data) on R (70). Protein annotations were characterized using DAVID as above. A heat map of differentially-pulled proteins was generated with the NMF package.

### Statistical analysis

Values are presented as mean±SE. A 2-tailed Student’s *t*-test was used to assess differences among groups. A *p* value of less than 0.05 was considered significant.

## Acknowledgements

We thank R. Gherzi for KSRP (Genova, Italy) plasmids; K.P. Lu (Harvard Boston) for the Pin1 plasmid; the Smoler Proteomics Center (Haifa Technion, Haifa, Israel) and the De Botton Protein Profiling institute of the Nancy and Stephen Grand Israel National Center for Personalized Medicine (Weizmann Institute of Science, Rehovot, Israel) for the MS analysis; G.W. Vainer (Hadassah Hebrew University Medical Center) for helpful discussions and E. Piontek, for professional technical assistance in paraffin embedding and tissue sectioning. This work was supported by grants from the Israel Science Foundation (to T.N-M., ISF 642/16) and the US-Israel Binational Science Foundation (to T.N-M. and I.Z.B-D., BSF 2019300). T.N-M. and M.N. are research associates of the Wohl’s Translation Research Institute at Hadassah-Hebrew University Medical Center. N.L. is the incumbent of the Alan and Laraine Fischer Career Development Chair and is supported by the Estate of Emile Mimran, Honey and Dr. Barry Sherman Lab, Dr. Barry Sherman Institute for Medicinal Chemistry, and Nelson P. Sirotsky.

