## Supplementary material for "Kidney failure alters parathyroid Pin1 phosphorylation and parathyroid hormone mRNA binding proteins leading to secondary hyperparathyroidism": Figures

### Slide 1
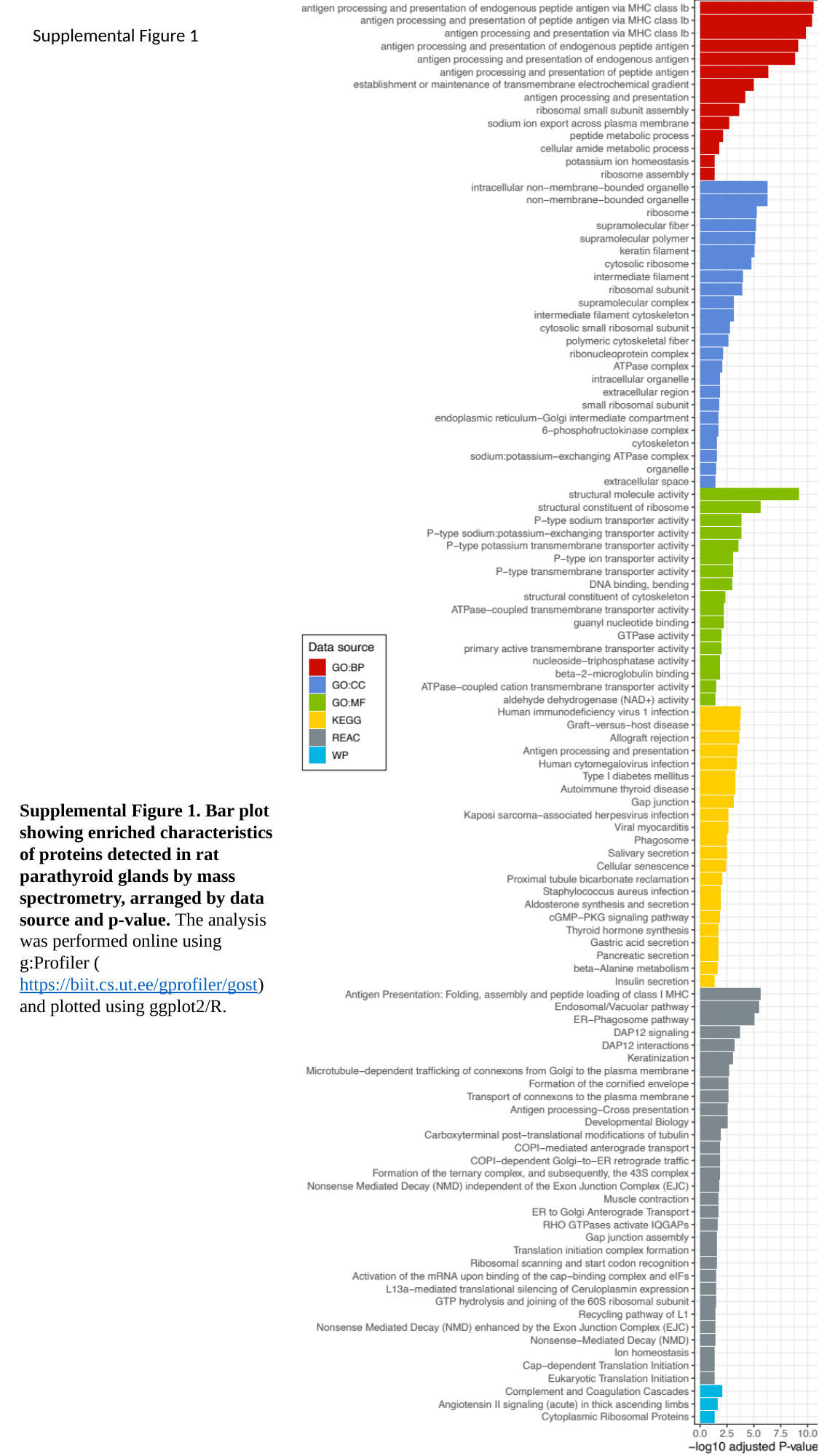

Supplemental Figure 1
Supplemental Figure 1. Bar plot showing enriched characteristics of proteins detected in rat parathyroid glands by mass spectrometry, arranged by data source and p-value. The analysis was performed online using g:Profiler (https://biit.cs.ut.ee/gprofiler/gost) and plotted using ggplot2/R.

### Slide 2
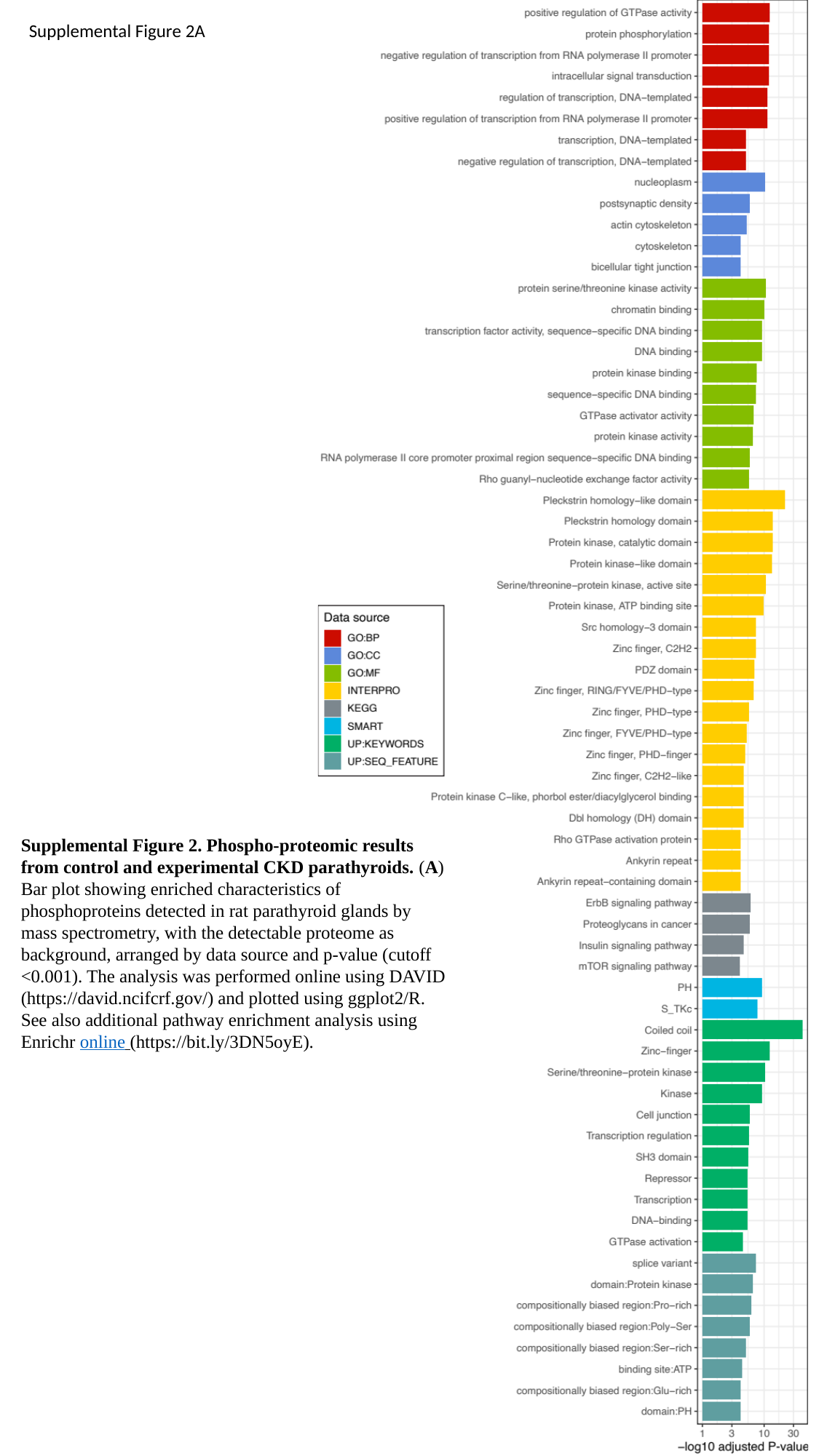

Supplemental Figure 2A
Supplemental Figure 2. Phospho-proteomic results from control and experimental CKD parathyroids. (A) Bar plot showing enriched characteristics of phosphoproteins detected in rat parathyroid glands by mass spectrometry, with the detectable proteome as background, arranged by data source and p-value (cutoff <0.001). The analysis was performed online using DAVID (https://david.ncifcrf.gov/) and plotted using ggplot2/R. See also additional pathway enrichment analysis using Enrichr online (https://bit.ly/3DN5oyE).

### Slide 3
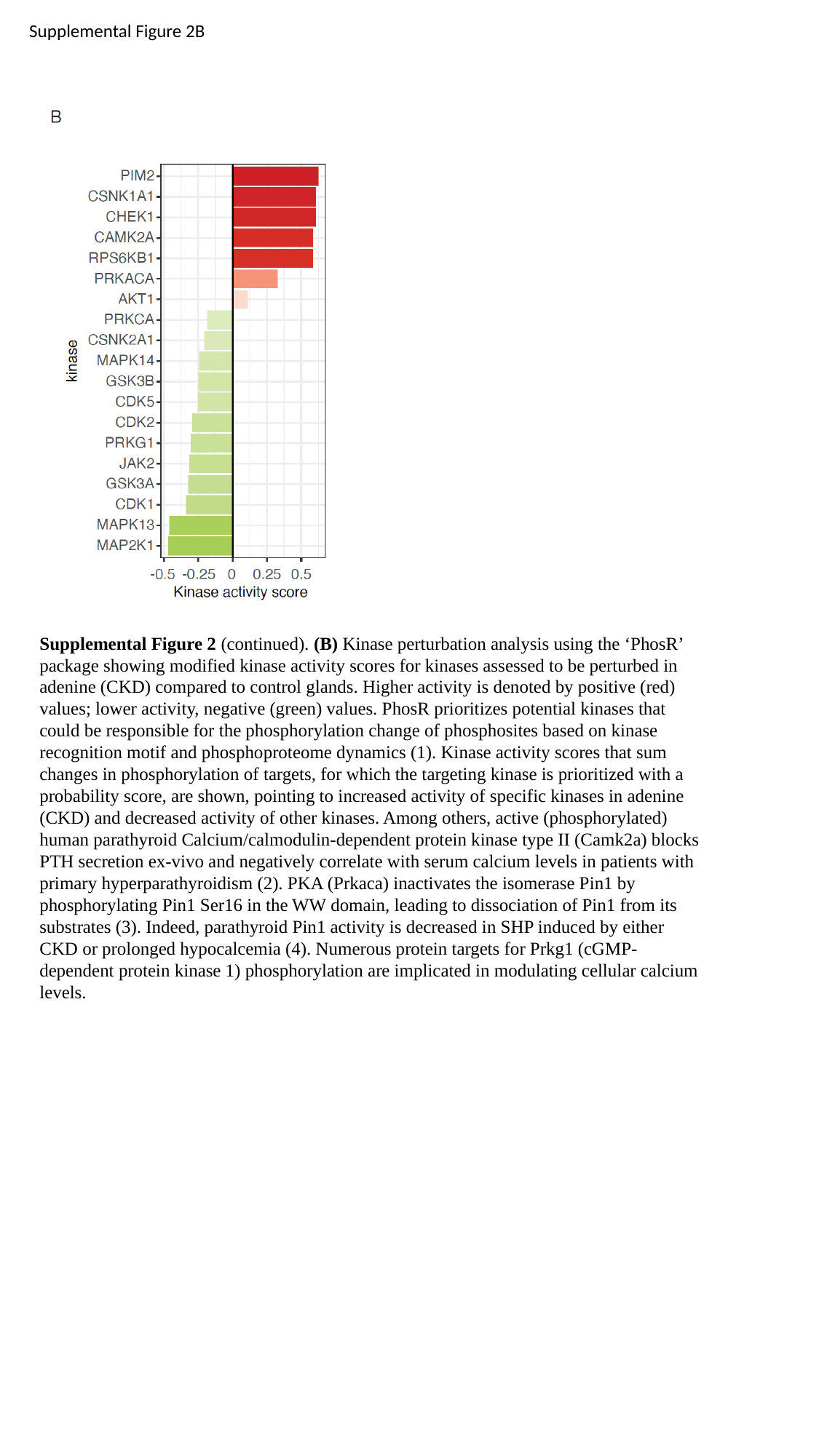

Supplemental Figure 2B
Supplemental Figure 2 (continued). (B) Kinase perturbation analysis using the ‘PhosR’ package showing modified kinase activity scores for kinases assessed to be perturbed in adenine (CKD) compared to control glands. Higher activity is denoted by positive (red) values; lower activity, negative (green) values. PhosR prioritizes potential kinases that could be responsible for the phospho­rylation change of phosphosites based on kinase recognition motif and phospho­proteome dynamics (1). Kinase activity scores that sum changes in phosphorylation of targets, for which the targeting kinase is prioritized with a probability score, are shown, pointing to increased activity of specific kinases in adenine (CKD) and decreased activity of other kinases. Among others, active (phosphorylated) human parathyroid Calcium/calmodulin-dependent protein kinase type II (Camk2a) blocks PTH secretion ex-vivo and negatively correlate with serum calcium levels in patients with primary hyperpara­thyroidism (2). PKA (Prkaca) inactivates the isomerase Pin1 by phosphorylating Pin1 Ser16 in the WW domain, leading to dissociation of Pin1 from its substrates (3). Indeed, parathyroid Pin1 activity is decreased in SHP induced by either CKD or prolonged hypocalcemia (4). Numerous protein targets for Prkg1 (cGMP-dependent protein kinase 1) phosphorylation are implicated in modulating cellular calcium levels.

### Slide 4
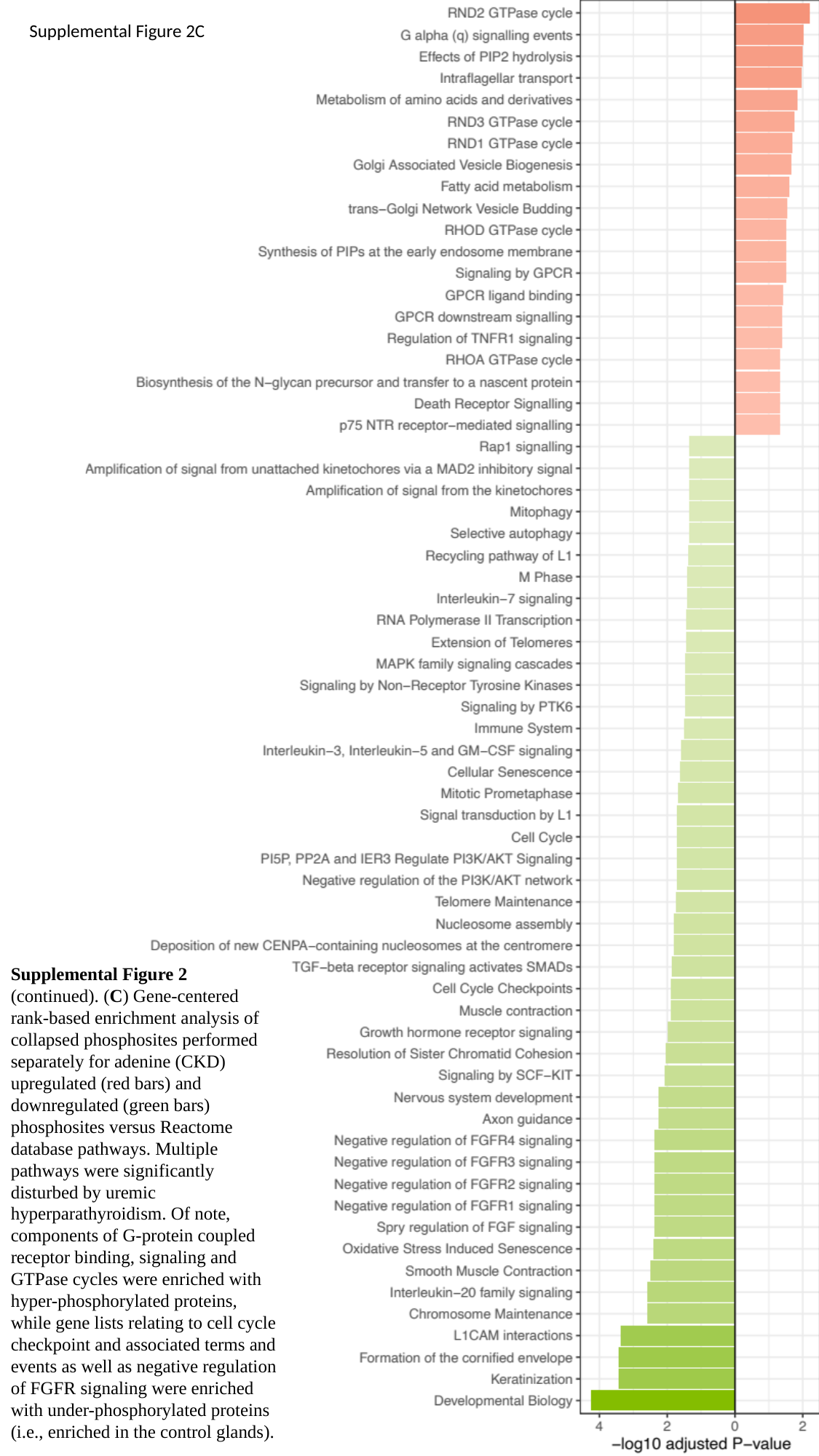

Supplemental Figure 2C
Supplemental Figure 2 (continued). (C) Gene-centered rank-based enrichment analysis of collapsed phosphosites performed separately for adenine (CKD) upregulated (red bars) and downregulated (green bars) phosphosites versus Reactome database pathways. Multiple pathways were significantly disturbed by uremic hyperparathyroidism. Of note, components of G-protein coupled receptor binding, signaling and GTPase cycles were enriched with hyper-phosphorylated proteins, while gene lists relating to cell cycle checkpoint and associated terms and events as well as negative regulation of FGFR signaling were enriched with under-phosphorylated proteins (i.e., enriched in the control glands).

### Slide 5
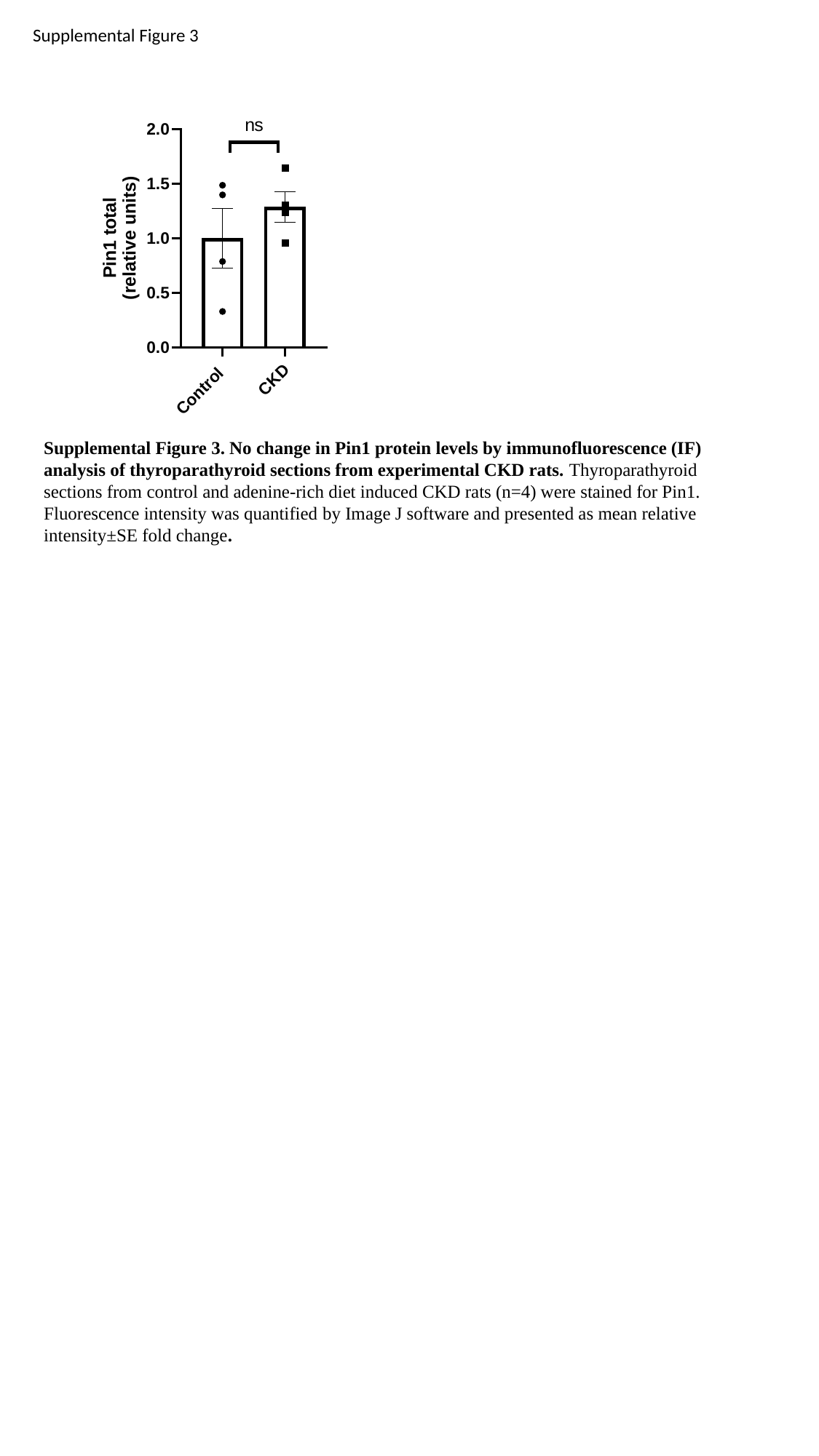

Supplemental Figure 3
Supplemental Figure 3. No change in Pin1 protein levels by immunofluorescence (IF) analysis of thyroparathyroid sections from experimental CKD rats. Thyroparathyroid sections from control and adenine-rich diet induced CKD rats (n=4) were stained for Pin1. Fluorescence intensity was quantified by Image J software and presented as mean relative intensity±SE fold change.

### Slide 6
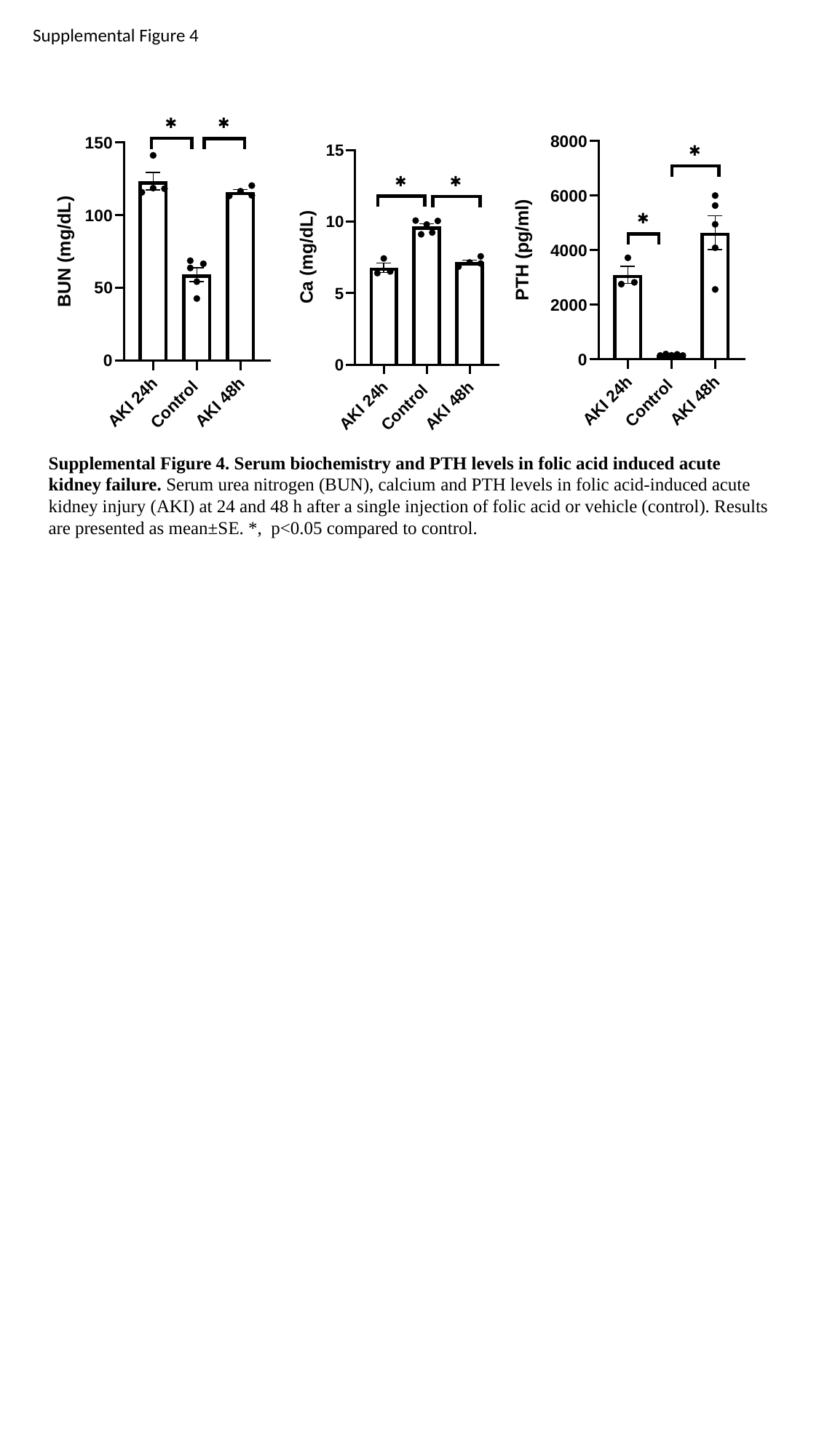

Supplemental Figure 4
Supplemental Figure 4. Serum biochemistry and PTH levels in folic acid induced acute kidney failure. Serum urea nitrogen (BUN), calcium and PTH levels in folic acid-induced acute kidney injury (AKI) at 24 and 48 h after a single injection of folic acid or vehicle (control). Results are presented as mean±SE. *, p<0.05 compared to control.

### Slide 7
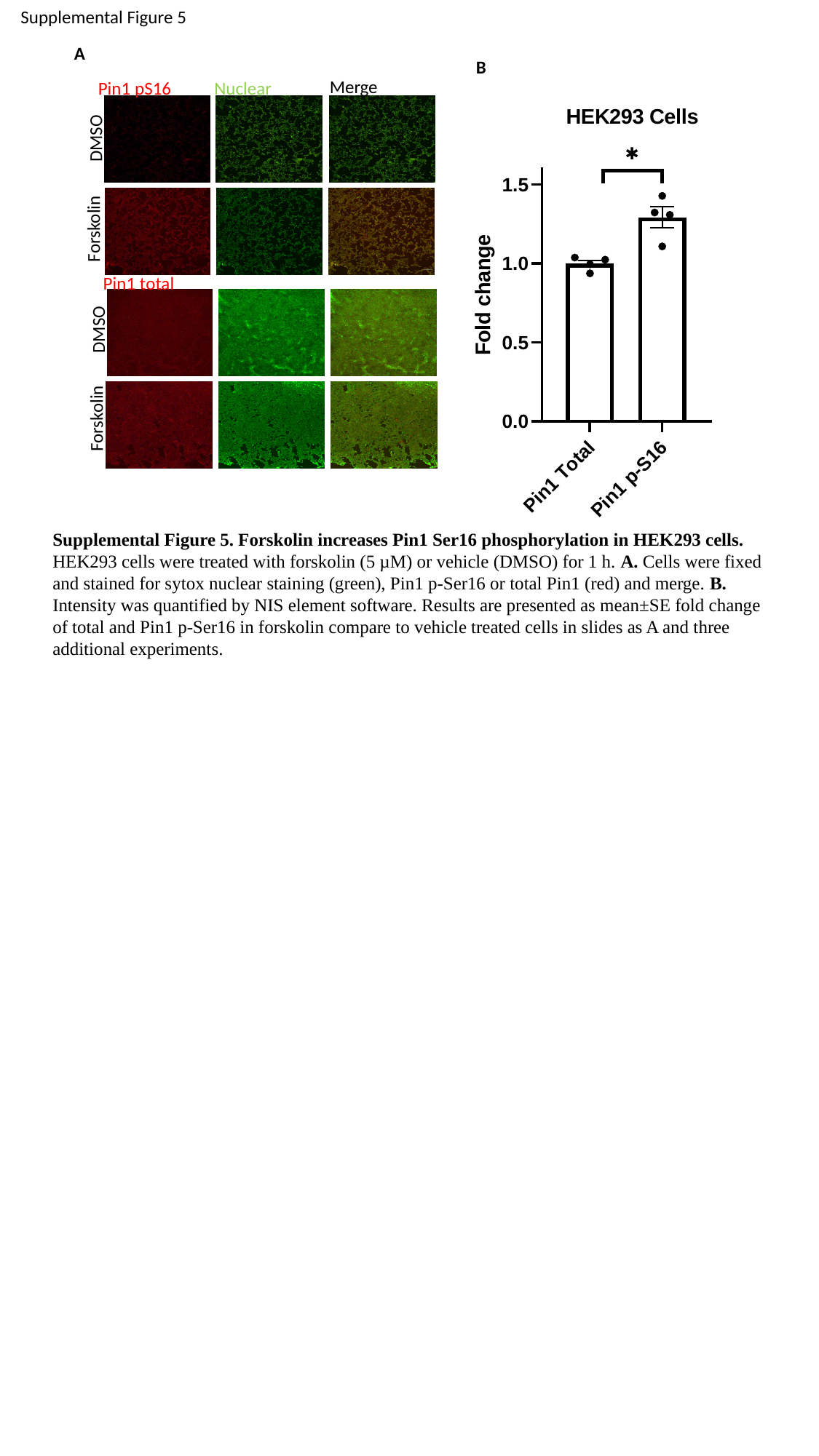

Supplemental Figure 5
A
B
Merge
Pin1 pS16
Nuclear
DMSO
Forskolin
Pin1 total
DMSO
Forskolin
Supplemental Figure 5. Forskolin increases Pin1 Ser16 phosphorylation in HEK293 cells. HEK293 cells were treated with forskolin (5 µM) or vehicle (DMSO) for 1 h. A. Cells were fixed and stained for sytox nuclear staining (green), Pin1 p-Ser16 or total Pin1 (red) and merge. B. Intensity was quantified by NIS element software. Results are presented as mean±SE fold change of total and Pin1 p-Ser16 in forskolin compare to vehicle treated cells in slides as A and three additional experiments.

### Slide 8
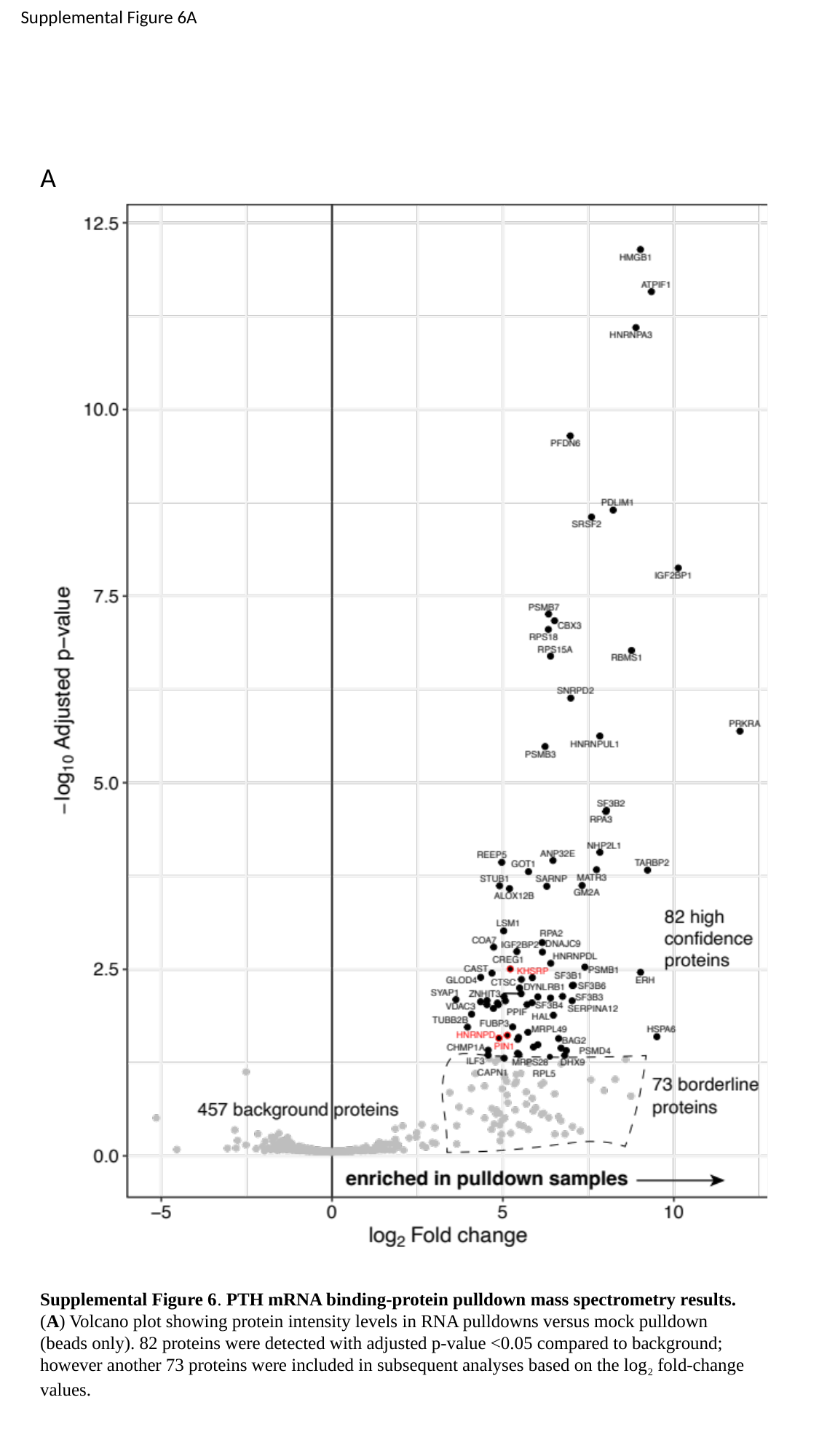

Supplemental Figure 6A
A
Supplemental Figure 6. PTH mRNA binding-protein pulldown mass spectrometry results. (A) Volcano plot showing protein intensity levels in RNA pulldowns versus mock pulldown (beads only). 82 proteins were detected with adjusted p-value <0.05 compared to background; however another 73 proteins were included in subsequent analyses based on the log_2_ fold-change values.

### Slide 9
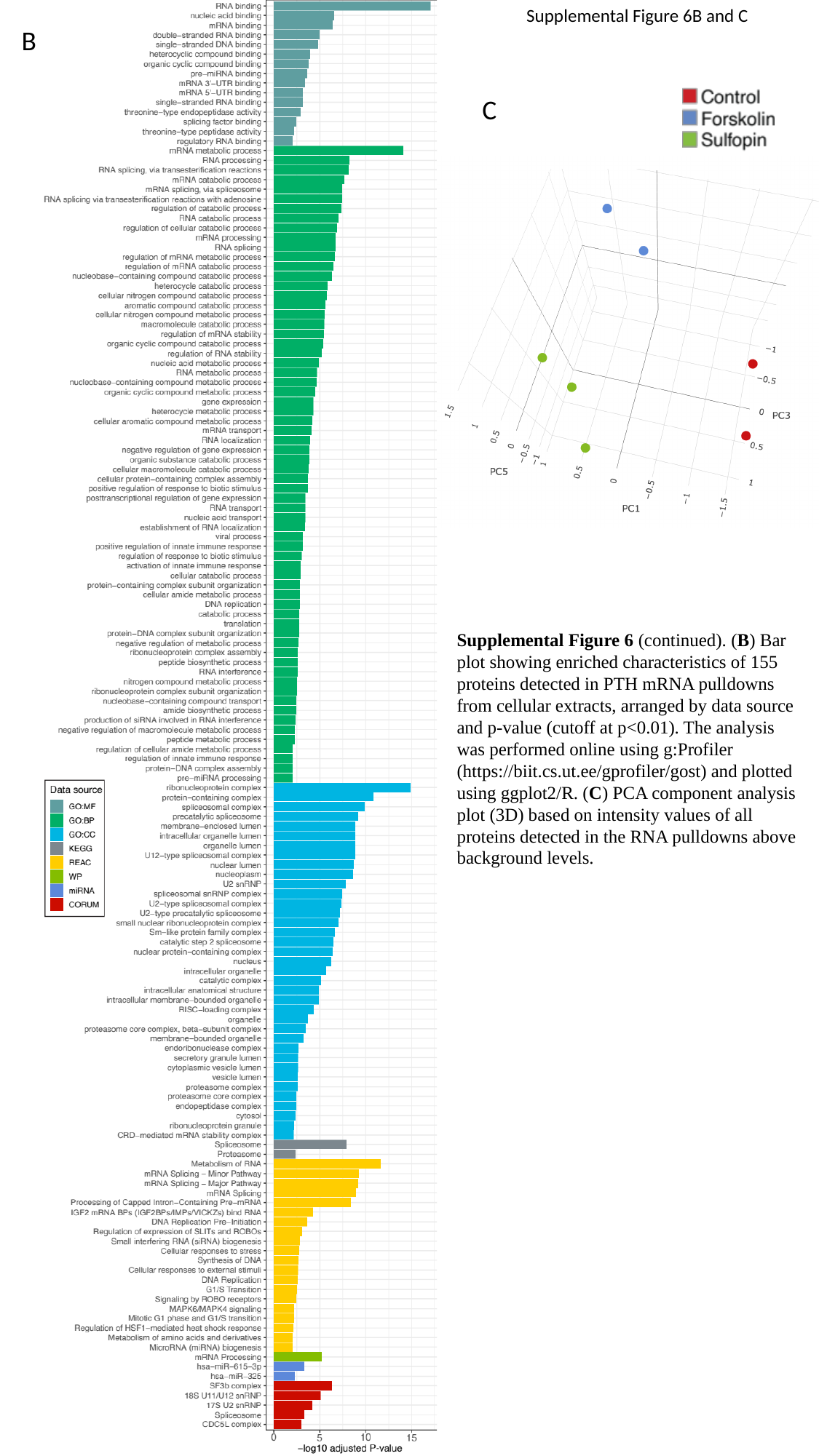

Supplemental Figure 6B and C
B
C
Supplemental Figure 6 (continued). (B) Bar plot showing enriched characteristics of 155 proteins detected in PTH mRNA pulldowns from cellular extracts, arranged by data source and p-value (cutoff at p<0.01). The analysis was performed online using g:Profiler (https://biit.cs.ut.ee/gprofiler/gost) and plotted using ggplot2/R. (C) PCA component analysis plot (3D) based on intensity values of all proteins detected in the RNA pulldowns above background levels.

### Slide 10
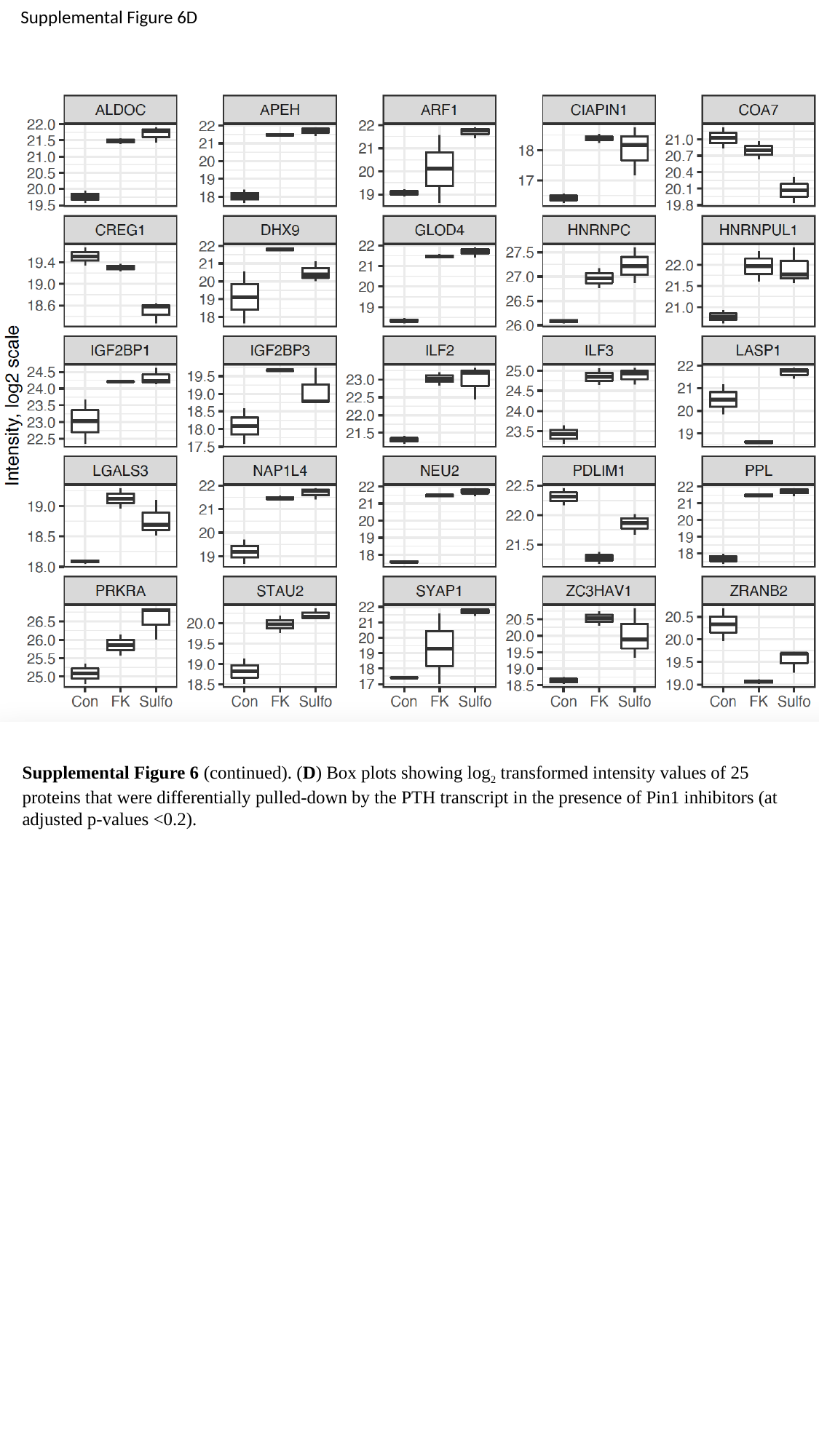

Supplemental Figure 6D
Supplemental Figure 6 (continued). (D) Box plots showing log_2_ transformed intensity values of 25 proteins that were differentially pulled-down by the PTH transcript in the presence of Pin1 inhibitors (at adjusted p-values <0.2).

### Slide 11
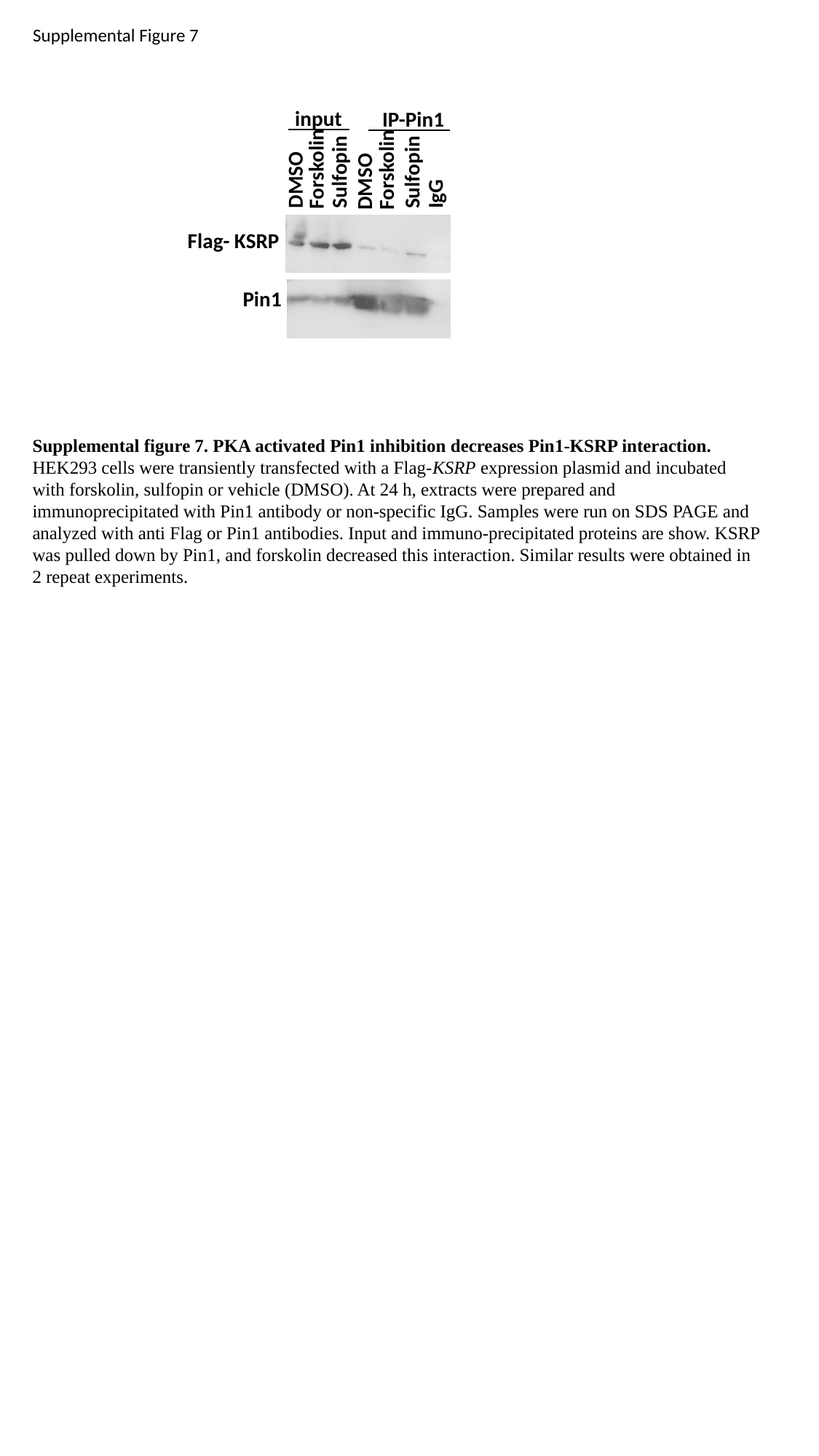

Supplemental Figure 7
DMSO
Sulfopin
Sulfopin
IgG
input
IP-Pin1
Forskolin
Forskolin
DMSO
Flag- KSRP
Pin1
Supplemental figure 7. PKA activated Pin1 inhibition decreases Pin1-KSRP interaction. HEK293 cells were transiently transfected with a Flag-KSRP expression plasmid and incubated with forskolin, sulfopin or vehicle (DMSO). At 24 h, extracts were prepared and immunoprecipitated with Pin1 antibody or non-specific IgG. Samples were run on SDS PAGE and analyzed with anti Flag or Pin1 antibodies. Input and immuno-precipitated proteins are show. KSRP was pulled down by Pin1, and forskolin decreased this interaction. Similar results were obtained in 2 repeat experiments.

### Slide 12
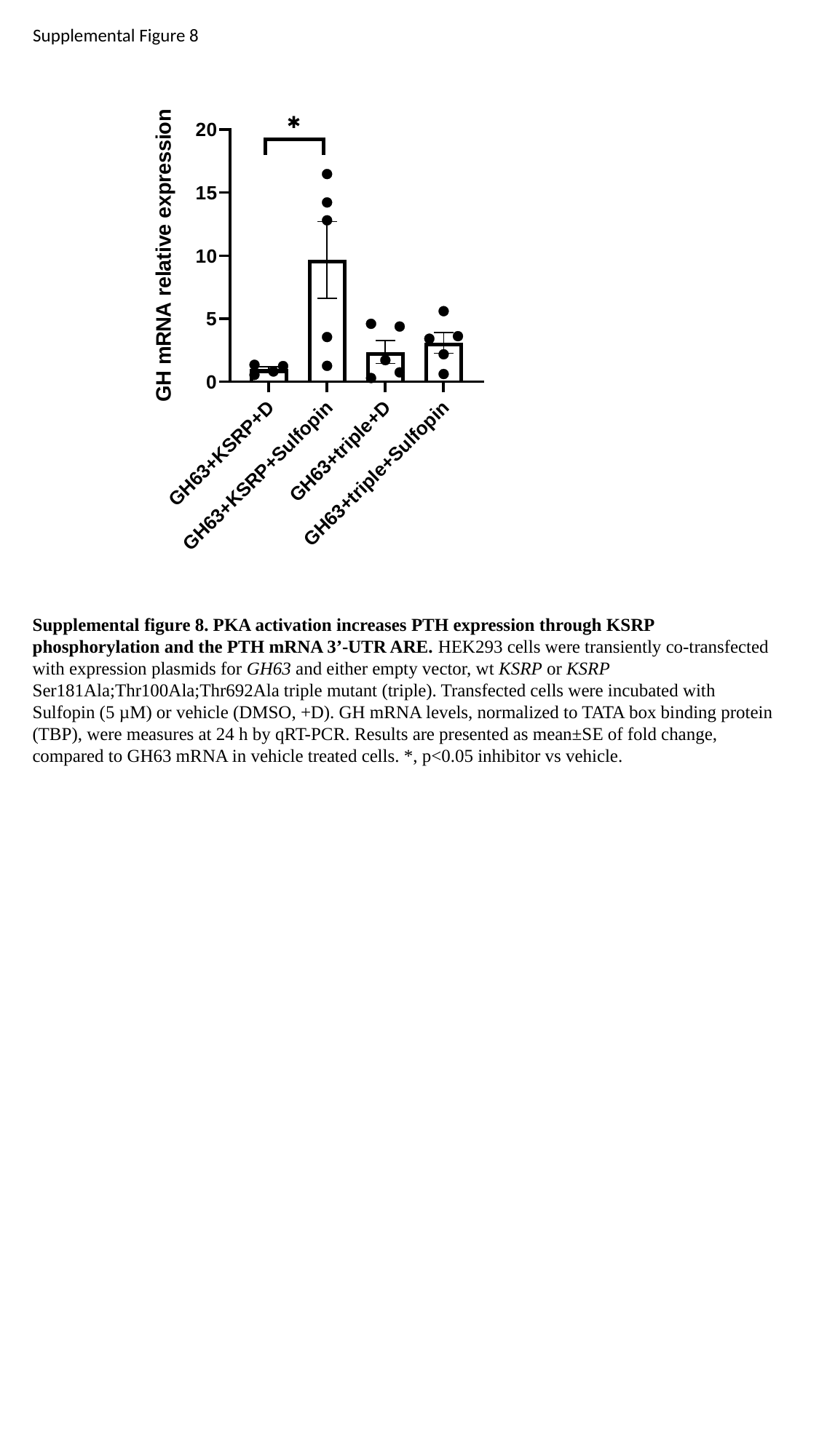

Supplemental Figure 8
Supplemental figure 8. PKA activation increases PTH expression through KSRP phosphorylation and the PTH mRNA 3’-UTR ARE. HEK293 cells were transiently co-transfected with expression plasmids for GH63 and either empty vector, wt KSRP or KSRP Ser181Ala;Thr100Ala;Thr692Ala triple mutant (triple). Transfected cells were incubated with Sulfopin (5 µM) or vehicle (DMSO, +D). GH mRNA levels, normalized to TATA box binding protein (TBP), were measures at 24 h by qRT-PCR. Results are presented as mean±SE of fold change, compared to GH63 mRNA in vehicle treated cells. *, p<0.05 inhibitor vs vehicle.
